# Screening of small molecule chemical library identifies leads that reduce cholesterol accumulation in human NPC1 fibroblasts

**DOI:** 10.1101/2022.11.16.516777

**Authors:** Nina H. Pipalia, Amy Y. Huang, Harold Ralph, Frederick R. Maxfield

## Abstract

Although considerable progress has been made in finding a therapy for Niemann-Pick type C1 (NPC1) disease, there still is no effective approved treatment. Previously, we reported results from a small molecule screen that was conducted on Chinese Hamster Ovary (CHO) cells (CT60) that lacked NPC1 expression. The hit compounds that were effective in reducing cholesterol accumulation in CT60 cells were not effective on human cells bearing disease-causing mutations in NPC1. Hence, we conducted a new small molecule screen on patient-derived NPC1 skin fibroblasts using a combinatorial library of ~48,000 compounds. We have identified several compounds that corrected the phenotype in human heterozygous mutant NPC1 skin fibroblasts (GM03123) bearing the inactivating I1061T mutation in one allele and another set of inactivating mutations in the other. Among the hit compounds were several histone deacetylase inhibitors, which we have explored further and reported previously. With the determination of the structure of NPC1 protein and the development of computational docking algorithms, it seems worthwhile to show the structures of 128 compounds that reproducibly reduced cholesterol storage in mutant NPC1 fibroblasts but with no known mechanism of action. These compounds were not toxic and specifically effective on mutant cells but not on NPC1^−/−^ cells, indicating on-target effect directly or indirectly.

## Introduction

Niemann Pick Type C1 (NPC1) disease is an often, fatal neurodegenerative lysosomal storage disorder most frequently caused by mutations in the *NPC1* gene that encodes for a late endosomal-lysosomal (LE/Ly) resident transmembrane protein, NPC1 (1). NPC1 disease is characterized by abnormal accumulation of cholesterol and glycosphingolipids, including GM2 and GM3 gangliosides, in the LE/Ly compartments of cells in multiple tissues (2, 3). Clinical manifestations include hepatosplenomegaly and progressive neurodegeneration that often leads to death in adolescence (4, 5). However, several late onset cases in adults with the same disease-associated mutations have been reported (6).

Structural and biochemical studies are elucidating the mechanism by which NPC1 protein promotes cholesterol efflux from LE/Ly. Cholesteryl esters are delivered to LE/Ly by endocytosis of lipoproteins (7), and these esters are hydrolyzed by lysosomal acid lipase (7). The cholesterol accumulates initially in small internal membranes inside the LE/Ly, and it is transferred from these membranes to a small soluble protein, NPC2 (8). NPC2 then delivers cholesterol to the N-terminal domain of NPC1, which is a multi-span protein in the limiting membrane of the LE/Ly (9). It appears that cholesterol can then move along a hydrophobic channel in NPC1 and be delivered to the membrane of the LE/Ly (10, 11), where it can then be delivered to other organelles by sterol transport proteins (12–15).

Therapies using chemical chelators of cholesterol such as 2-hydroxylpropyl beta-cyclodextrin that bypass the need for NPC1 or NPC2 (16, 17) were found to be effective in reversing the defect in mouse and cat models of NPC1 (18). A human clinical trial of cyclodextrin and continued compassionate use are in progress, but central nervous system therapy requires direct injection into the cerebrospinal fluid since cyclodextrins do not cross the blood brain barrier efficiently (19, 20).

As described in this paper, we conducted a screen of ~ 48,000 small molecules on human NPC1 mutant skin fibroblasts (GM03123), which were from a compound heterozygote patient. One allele carried a pathogenic splice site mutation, c.1947+5G>C as well as P237S, and the second allele carried a pathogenic missense mutation, I106T. We identified 129 compounds that reproducibly reduced the NPC1 cholesterol storage phenotype at concentrations at or below10 µM, with several compounds effective at lower concentrations. We found that several histone deacetylase inhibitors (HDACi) were effective in reversing the NPC1 mutant phenotype, and they led to increased expression of the NPC1 protein and its correct delivery to LE/Ly (21). However, in a study in a mouse model of NPC1^I1061T^, it was found that even HDACi with fairly good brain penetration could not rescue the NPC1 defects in an animal (22). Nevertheless, the HDACi studies did show that, for the mutants tested, we could rescue the NPC cholesterol storage phenotype by facilitating proper folding and transport of the mutant NPC1 proteins (21). A subsequent analysis of 60 human mutant NPC1 proteins expressed in NPC1 null U20S cells showed that ~85% of the mutants could be rescued in an analysis of their cholesterol storage in LE/Ly (23). These studies indicated that most NPC1 human mutations could be rescued by treatments that facilitate folding of the mutant NPC1 proteins.

## Materials and Methods

Cell medium (MEM with Earle’s salts), L-glutamine and FBS were purchased from Invitrogen (Carlsbad, CA). The compound libraries for screening were purchased and maintained by the Weill Cornell/Rockefeller High Throughput Screening Resource Center (HTRSC) at The Rockefeller University. The chemicals were purchased from Chemical Diversity, Inc. (San Diego, CA), Prestwick Chemicals (France), Cerep (Seattle, WA), ChemBridge (San Diego, CA) and MicroSource Spectrum. Nuclear stain DRAQ5 was purchased from Biostatus Ltd. (UK). All other chemicals, including DMSO, filipin, and para-formaldehyde (PFA) were purchased from Sigma Chemicals (St. Louis, MO). MetaXpress image-analysis software was from Molecular Devices (Downington, PA).

### Compound libraries

The library used for the chemical screen contained 47,683 chemicals. Many of these are biologically active and structurally diverse compounds based on known drugs, experimental bioactives, and pure natural products. The ChemDiv library was a combinatorial library consisting of 125 combinatorial templates encompassing total 25,947 compounds. The ChemBridge library included 5,000 low molecular weight drug-like compounds. The Cerep library included 4,000 drug-like molecules based on 350 scaffolds. The in-house focused library included 9856 compounds. The MicroSource Spectrum collection included 2000 drug components, natural products and bioactive compounds. The Prestwick chemical library was a collection of 880 high purity chemical compounds of which 85% are marketed drugs, and 10% are bioactive alkaloids or related substances.

### Cell Culture and Media

Wild type (WT) normal human fibroblasts (GM5659) and human Niemann-Pick Disease, type C1 (NPC1) fibroblasts (GM03123). GM03123 fibroblasts were from a compound heterozygote patient; one allele carried a pathogenic splice site mutation c.1947+5G>C as well as P237S and the second allele carried a pathogenic missense mutation I106T. They were purchased from Coriell Cell Repositories (Camden, NJ) and were grown in Medium A consisting of Minimum essential medium (MEM) with Earle’s salts, 2 mM L-glutamine and 10% FBS. Screening Medium S was prepared by supplementing Medium A with 20 mM HEPES. The concentration of the primary stock of tested chemicals in the library was 5 mM in DMSO.

### Screening Protocol

Cells were plated in 384 well tissue culture treated plates (Corning Inc, NY) at 600-650 cells per well in 30 µl growth medium (MEM + 10% FBS medium) using Multidrop (Thermo Fisher Scientific, Inc.). NPC1 fibroblasts were plated in the first 23 columns on a 384-well plate. Normal fibroblasts were plated in column 24 and used for the determination of Z’ values. Immediately after plating, cells were left at room temperature for 30 min to settle and then incubated in a CO2 incubator at 37° C. After 24 hours, compound mixtures were added at a final concentration of 10 µM. A 2X stock of compounds was prepared by mixing 0.1 µl of a primary stock (5 mM stock in DMSO) in 25 µl of Medium S in Falcon 384-well V-bottom polypropylene plates using a Packard MiniTrak robotic liquid-handling system. To obtain final concentrations of 10 µM, 2X stock of compounds were dispensed into plates containing cells in growth medium. For primary and secondary screening, 352 test compounds were added to each plate (columns 1-22), out of the remaining 32 wells column 23 was used as a solvent control by adding only DMSO, and in column 24 WT fibroblasts were plated as a negative control. All plates were incubated with compounds for 16-20 hrs at 37°C. Plates were then washed three times with phosphate buffer saline (PBS), pH 7.4, using a Bio-Tek Elx405 plate washer (Bio-Tek Instruments, Inc., Winooski, VT). For each wash cycle, 70 µl of PBS was dispensed followed by aspiration with a residual volume of < 16 µl/well. The cells were fixed with 1.5 % PFA in PBS for 30 min at room temperature, with gentle shaking on a Wellmix shaker (Thermo Scientific), followed by three more PBS washes.

### Fluorescence labeling

To the fixed cells, filipin was added at a final concentration of 50 µg/ml in PBS and incubated for 45 min at room temperature. Cells were then washed three times with PBS. To stain nuclei, DRAQ5 was added at a final concentration of 1.5 µM. Plates were imaged after overnight staining with DRAQ5.

### Fluorescence microscopy

An ImageXpress^MICRO^ imaging system (Molecular Devices) equipped with a 300W Xenon-arc lamp from Perkin-Elmer, a Nikon 10X Plan Fluor 0.3 NA objective, and a Photometrics CoolSnapHQ camera (1,392 × 1,040 pixels) from Roper Scientific was used to acquire images. Filipin images were acquired using 377/50 nm excitation and 447/60 nm emission filters with a 415 nm dichroic long-pass filter. DRAQ5 images were acquired using 628/40 nm excitation and 692/40 nm emission filters with a 669 nm dichroic long-pass filter. Filter sets were obtained from Semrock.

Plates were transported from plate hotels using a CRS CataLyst Express robot from Thermo Fisher Scientific. Images were acquired at four sites per well, each ~50 µm from the center of the well, with 25 ms exposure time per image for filipin and 100 ms per image for DRAQ5 using 2 × 2 binning acquisition. Each site was individually focused using a high-speed laser autofocus comprised of a 690 nm diode laser and a dedicated 8-bit CMOS camera. Acquisition time was 75 minutes per plate. 696 × 520 pixel images were acquired at 12 intensity bits per pixel. Each pixel is 1.29 × 1.29 µm in the object.

### Image analysis

Images were analyzed using MetaXpress image-analysis software (Molecular Devices). We obtained three different values from the images: 1) Average filipin intensity in cells, 2) Lysosomal storage organelle (LSO) compartment ratio assay and 3) Cell count. The LSO compartment ratios, which are calculated as the total intensity above a defined high threshold divided by the total cell area, were estimated as described in detail previously (24). The high threshold was selected to identify the bright area of flipin fluorescence near the cell center, which contains the cholesterol-loaded LE/Ly. Cell area was defined as the area containing filipin fluorescence from other cell membranes. The threshold values for each set of dishes were chosen based on analysis of untreated NPC1 mutant cells. Normalized values were obtained by dividing the values in the presence of each compound by the average of the values obtained in the presence of the solvent control for each plate. A Z’ calculation (25) for each plate was done for the solvent controls (normal and NPC fibroblasts) to indicate plate and screen quality.

The cell count assay was done using the Count Nuclei Module in MetaXpress. Briefly, the approximate minimum nucleus width, the approximate maximum nucleus width and the intensity above background was set based on the DRAQ5 signal. Cell count number was used to identify toxic compounds.

## Results

### Primary Screen

The high content screen was performed using a previously validated fluorescence assay (24) using filipin labeling to visualize and quantify the amount of unesterified cholesterol in the whole cell as well as in LE/Ly compartments near the center of the cell. Filipin is a fluorescent polyene antibiotic that selectively binds to unesterified cholesterol. After 24h treatment, cells were fixed, stained, and imaged as described in detail in Methods. Shown in Figure 1A and 1B are representative filipin images of WT (GM5659) and NPC1 mutant (GM03123) skin fibroblasts, respectively. The Z’ value for the LSO compartment ratio used to analyze screening data for identification of hit compound was 0.55 as shown in Figure 1C. This indicated that the assay was robust enough to provide a fairly high degree of discrimination between WT and NPC1 fibroblasts (25). Using this high content screen, we examined 47,683 small molecule compounds from six different libraries of synthetic small molecules (Table 1). Generally, the total cell count for all four images per well ranged between 375-500 cells, which allowed us to look at the effects of the compounds on a large number of cells.

**Table 1.** Number of compounds tersted from different vendor libraries.

| **Vendor Library** | **Compounds** |
| --- | --- |
| Chemical Diversity 1 and 2 | 25947 |
| Prestwick | 880 |
| Cerep Odyssey II | 4000 |
| Focused Library (In house focused library) | 9856 |
| ChemBridge NovaCore | 5000 |
| Microsource Spectrum | 2000 |
| Total | 47683 |

**Figure 1.**
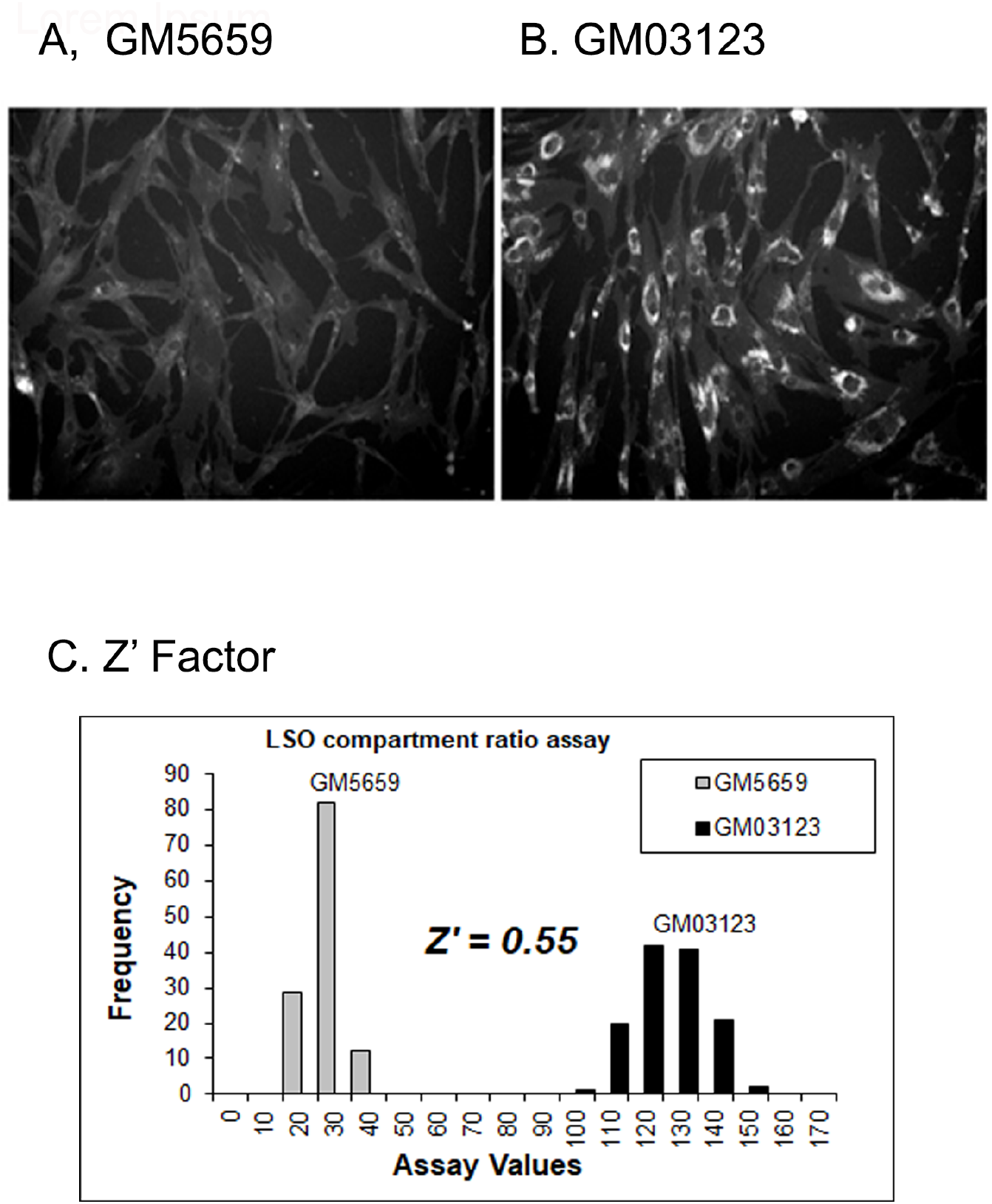
Representative images of filipin labeled cells. Normal Human fibroblasts (GM 5659) and fibroblasts (GM 03123) from a Niemann Pick disease type C patient were plated on 384 well plates and grown in growth medium for 48 h. Cells were washed 3 times with PBS, fixed with 1.5% PFA, and stained with filipin for 45 min at room temperature. Images were acquired using an ImageXpress ^MICRO^ automated microscope using a 10X objective using 377/50 nm excitation and 447/60 nm emission filters with a 415 dichroic long pass filter. (A) GM5659 normal human fibroblasts labeled with filipin. (B) NPC1 mutant (GM 03123) cells labeled with filipin. (C) Histogram of LSO ratio values for calculation of Z’ measure of screen robustness. X-axis represents raw LSO ratio assay value not normalized, and Y-axis represents frequency of occurrence

Images were corrected for shading and background intensity (24). We used a high threshold in the filipin images to identify the LE/Ly in the perinuclear region of NPC mutant cells (24). A low threshold value in the filipin image was used to identify the whole cell area. The detailed stepwise image analysis process for calculating lysosomal storage organelle (LSO) ratio is shown in Figure 2A. To compare data from different plates and different experiments, the LSO ratio values for each plate were normalized by dividing the LSO value of each image within a plate by that of an average LSO value of DMSO-treated NPC1 mutant cells (64 images from 16 wells of GM03123) in the corresponding plate. Thus, a normalized LSO value of 1.0 will be indicative of no effect (24). The standard deviation (SD) of DMSO-treated images in each plate was calculated, and only those compounds that showed an LSO ratio more than three SD values below the DMSO-treated mean were considered as a hit.

**Figure 2.**
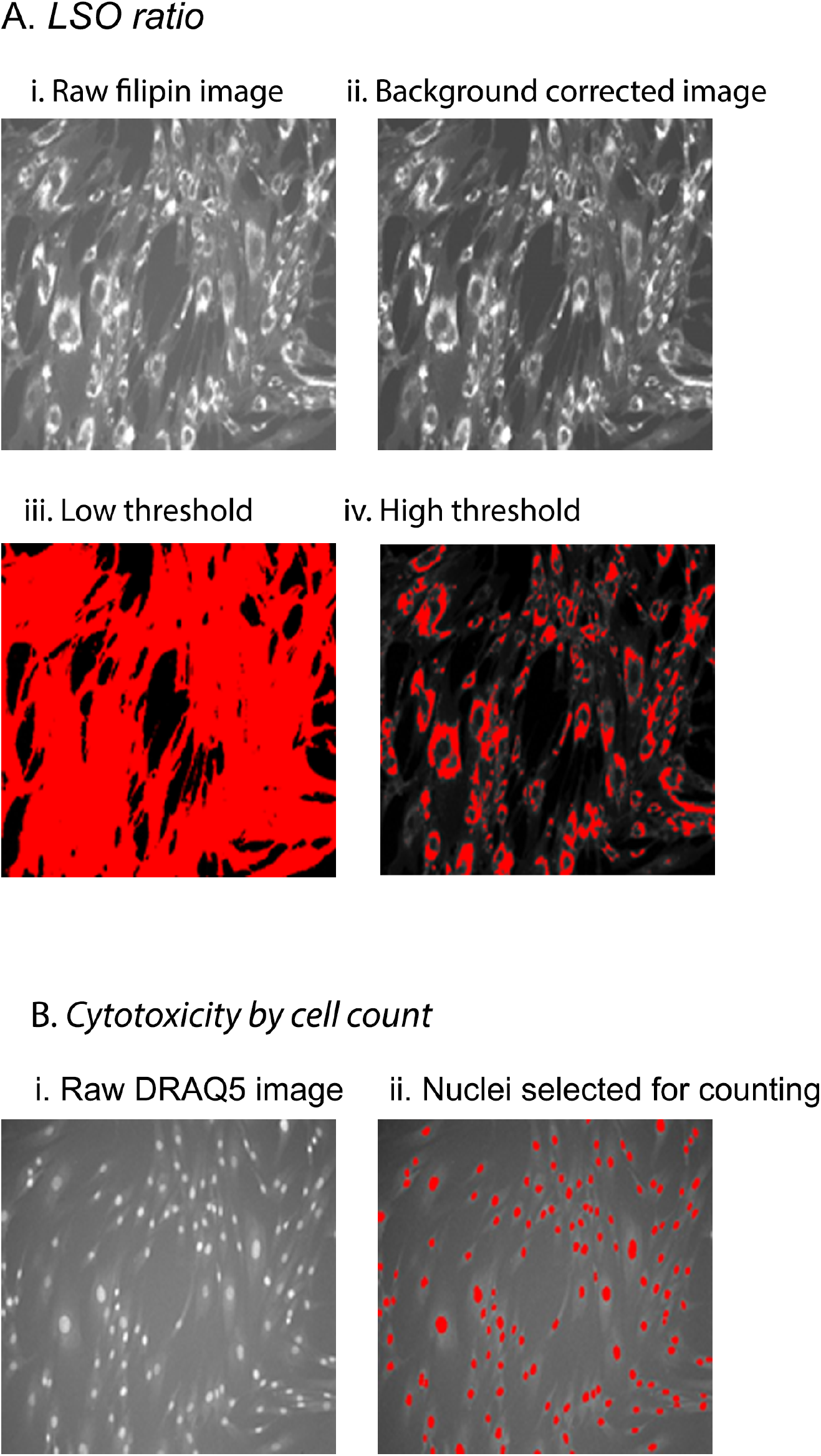
Illustration of image analysis methods. Cells were fixed with PFA and labeled with filipin and DRAQ5. Images were acquired using an ImageXpress ^MICRO^ automated fluorescence microscope with a 10X objective using 360/40 nm excitation and 480/40 nm emission filters with a 365 DCLP filter. LSO Ratio Measurement. (i) Raw filipin image, (ii) Shading and background-corrected image, (iii) A low-threshold image for identifying cell area, (iv) High threshold image used to identify LSO compartments. (B) Nuclear staining. (i) DRAQ5 stained image showing nuclei. (ii) Image showing objects identified as nuclei using the count nuclei application.

In order to eliminate cytotoxic compounds, we counted the number of cells in each image before and after treatment (Figure 2B). Any compound that caused greater than 25% cell loss was discarded. Also, out-of-focus images and wells with pipetting errors were marked and excluded from further consideration. Thus, 668 non-toxic hit compounds with significant reduction in LSO ratio were selected for further analysis, which is 1.9% of the total compounds screened. The primary screen was a single point determination, meaning each compound was added to only one well (Table 2).

**Table 2.** Compound elimination criteria and statistics. Shown in the table are the number of compounds eliminated from selection as a hit compound based on toxicity, technical errors associated with instruments, or other reasons like precipitation. ~800 compounds were eliminated, out of which 1.17% were due to imaging errors, including out of focus images, 0.83% of compounds were toxic as observed by reduced cell number, and less than 0.1% were due to pipetting errors or compound precipitation.

|  | No. of compounds | % Total |
| --- | --- | --- |
| Imaging error | 444 | 1.17% |
| Pipetting error | 39 | 0.10% |
| Precipitate | 3 | 0.01% |
| Toxic | 316 | 0.83% |

### Secondary Screen

*Cherry Picking Round One:* For confirmation, we selected the 668 hit compounds from the primary screen and repeated the assay two separate times. In each experiment, each compound was added in duplicate. Thus, each compound was tested at 10 µM in four different wells, and four images were acquired per well. These 16 values were averaged. The screening of 668 compounds validated 263 non-toxic compounds that were effective in reducing cholesterol accumulation after the first round of cherry picking which is 0.7 % of total compounds screened. They showed a strong positive effect (LSO reduction ≥3SD) in three out of four wells including 22 hits from the Prestwick library, which contains mainly FDA approved compounds.

*Dose Response Study of the hits from Secondary Screen:* The 263 hit compounds from the first round of cherry picking were subjected to a second round of screening at four different concentrations: 10, 1, 0.1 and 0.033 µM, in triplicate. Compound treatments that resulted in greater than 25% cell loss, as compared to DMSO-treated cells, were eliminated. Some of the hit compounds were eliminated based on previously reported information (e.g., toxicity) as explained in the Discussion.

After screening and evaluation of all hit compounds, the second round of cherry picking resulted in confirming 129 compounds that significantly reduced cholesterol accumulation in LSO compartments of NPC1 fibroblasts. Moreover, some of the hit compounds were effective at lower concentrations.

None of these 134 compounds were toxic to the cells, as observed by the cell count assay. A schematic in Figure 3 and Table 3 summarizes the normalized LSO index of all the 128 compounds at their respective doses. We observed that several compounds were effective in correcting NPC1 phenotype at all four concentrations tested (Table 3). We also noted that some compounds, including lysosomal acid lipase inhibitors, Orlistat and Lalistat 1, were effective at all four doses. These would not be effective therapeutically because reduced expression of lysosomal acid lipase is itself a serious genetic disorder, Wolman’s disease (26). We previously characterized the activity of these lipase inhibitors, and the results were published elsewhere (27–29).

**Table 3.**
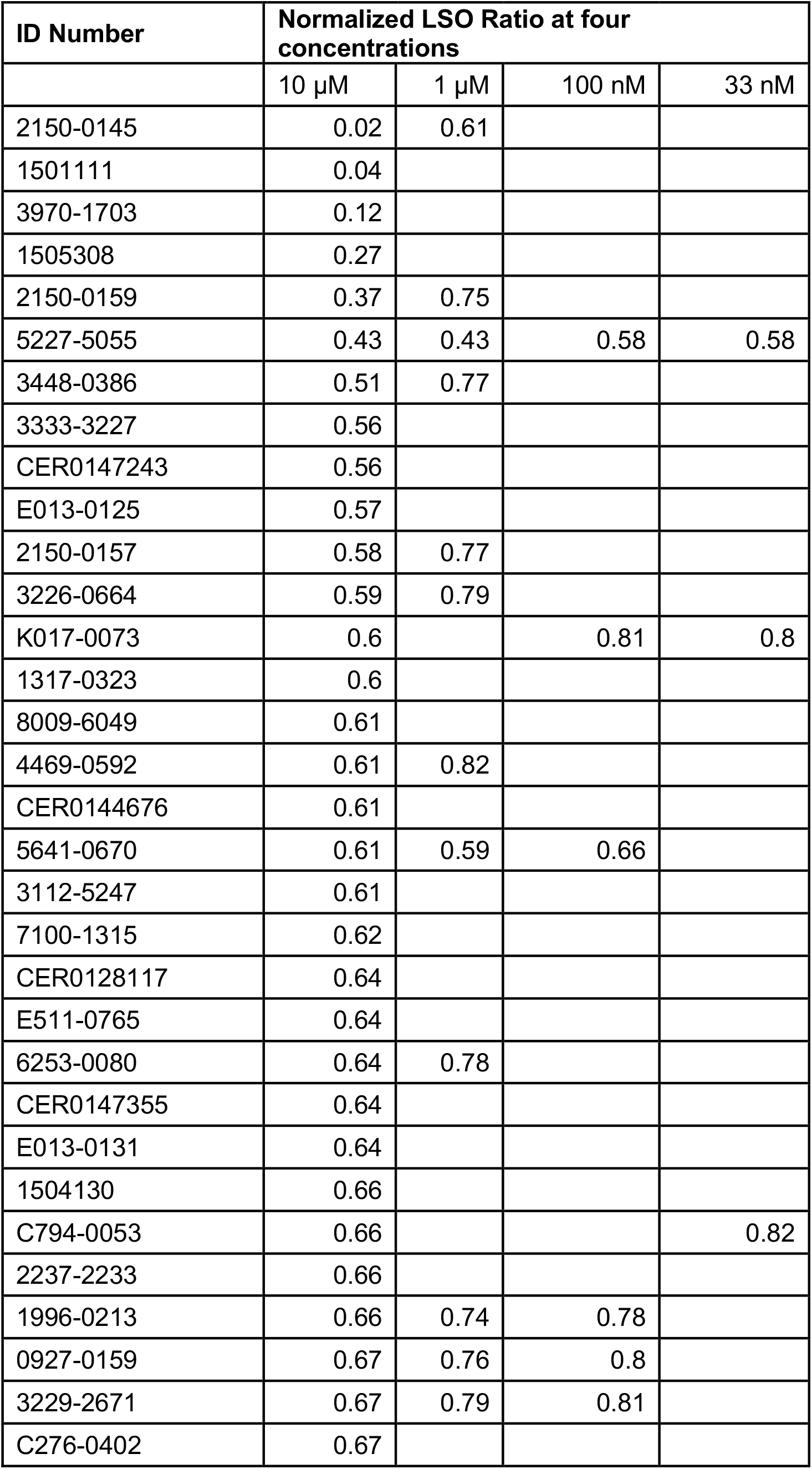
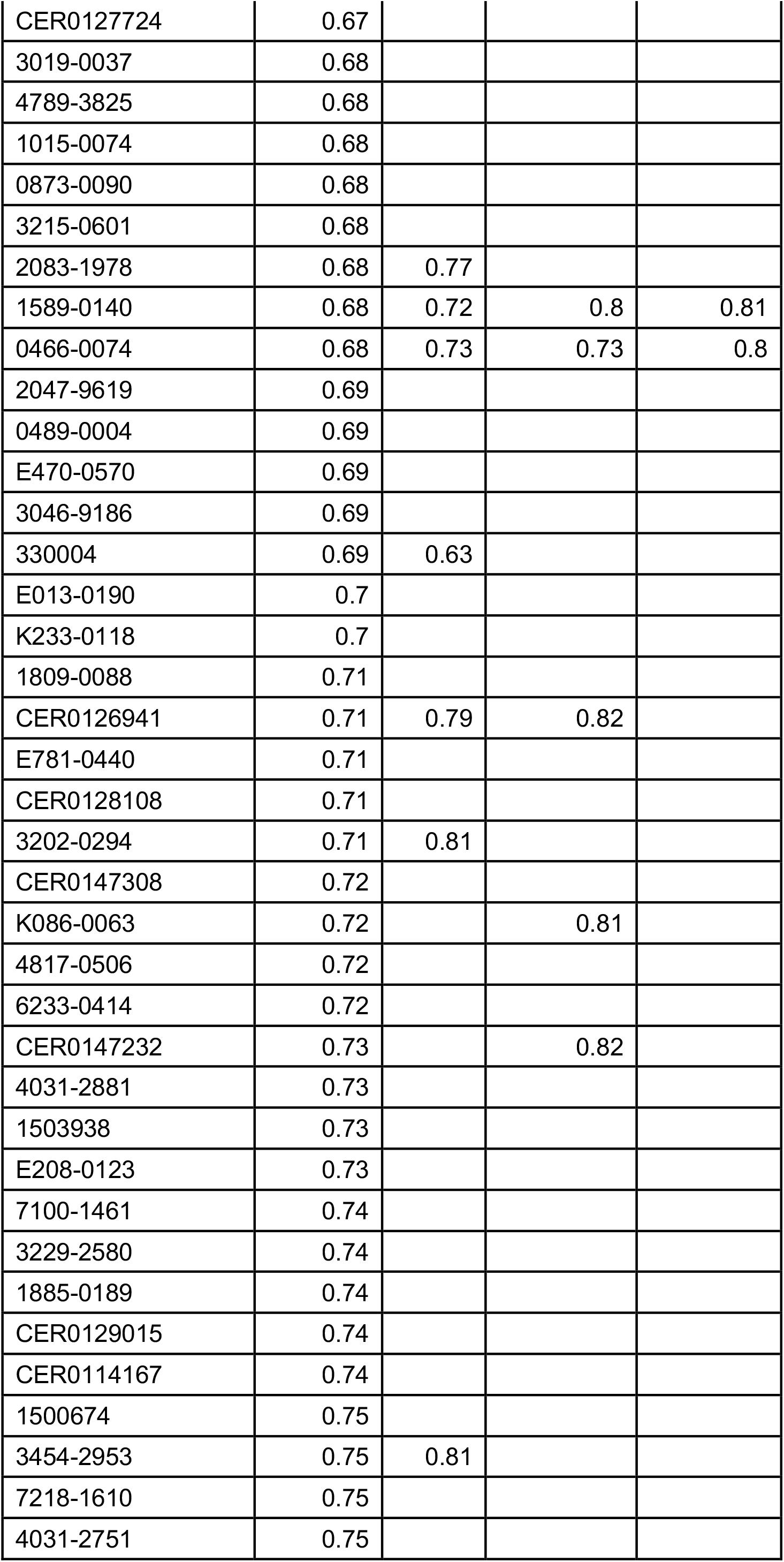
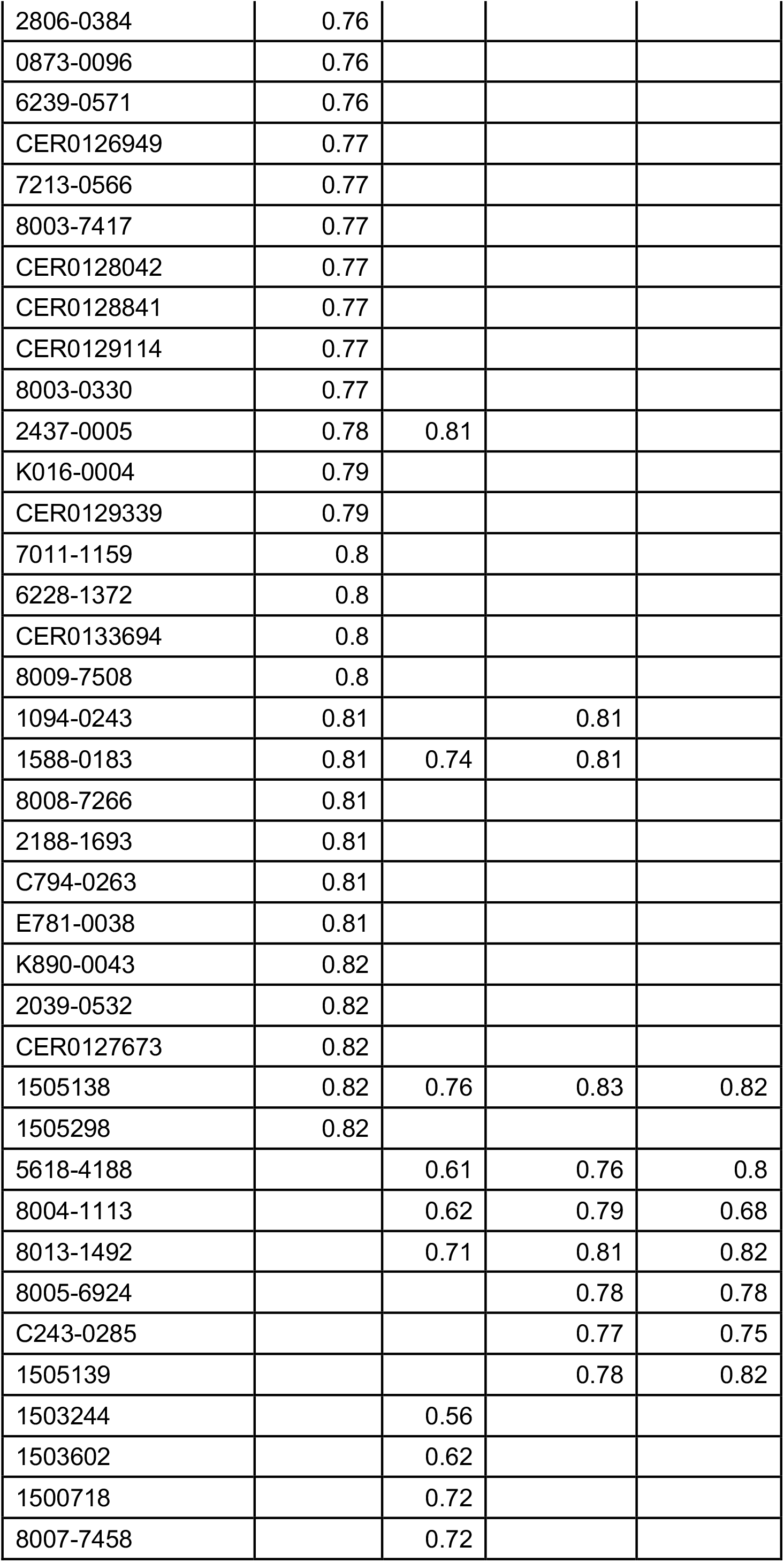
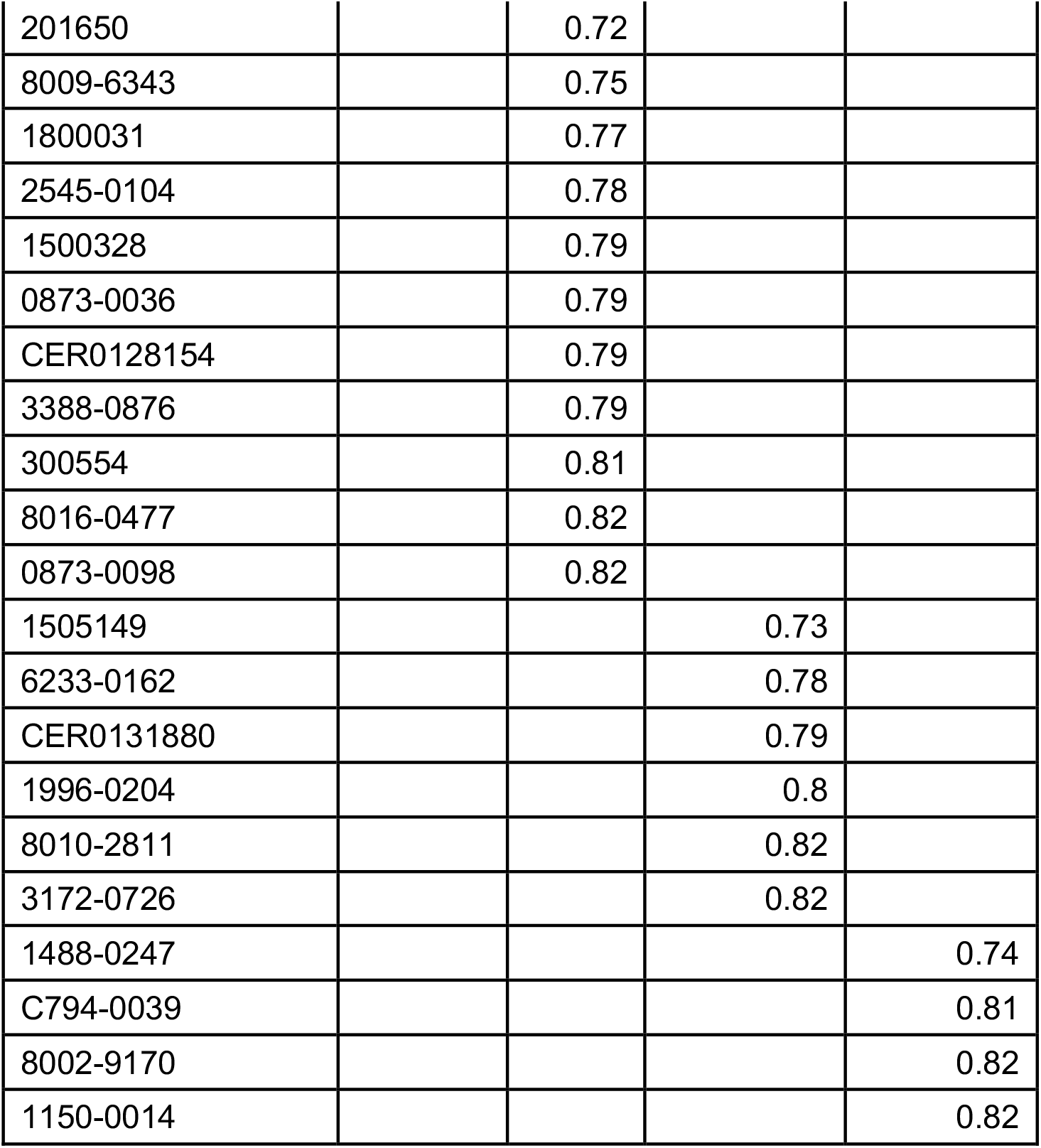
LSO values of hit compounds.

| ID Number | Normalized LSO Ratio at four concentrations |  |  |  |
| --- | --- | --- | --- | --- |
| | 10 μM | 1 μM | 100 nM | 33 nM |
| 2150-0145 | 0.02 | 0.61 |  |  |
| 1501111 | 0.04 |  |  |  |
| 3970-1703 | 0.12 |  |  |  |
| 1505308 | 0.27 |  |  |  |
| 2150-0159 | 0.37 | 0.75 |  |  |
| 5227-5055 | 0.43 | 0.43 | 0.58 | 0.58 |
| 3448-0386 | 0.51 | 0.77 |  |  |
| 3333-3227 | 0.56 |  |  |  |
| CER0147243 | 0.56 |  |  |  |
| E013-0125 | 0.57 |  |  |  |
| 2150-0157 | 0.58 | 0.77 |  |  |
| 3226-0664 | 0.59 | 0.79 |  |  |
| K017-0073 | 0.6 |  | 0.81 | 0.8 |
| 1317-0323 | 0.6 |  |  |  |
| 8009-6049 | 0.61 |  |  |  |
| 4469-0592 | 0.61 | 0.82 |  |  |
| CER0144676 | 0.61 |  |  |  |
| 5641-0670 | 0.61 | 0.59 | 0.66 |  |
| 3112-5247 | 0.61 |  |  |  |
| 7100-1315 | 0.62 |  |  |  |
| CER0128117 | 0.64 |  |  |  |
| E511-0765 | 0.64 |  |  |  |
| 6253-0080 | 0.64 | 0.78 |  |  |
| CER0147355 | 0.64 |  |  |  |
| E013-0131 | 0.64 |  |  |  |
| 1504130 | 0.66 |  |  |  |
| C794-0053 | 0.66 |  |  | 0.82 |
| 2237-2233 | 0.66 |  |  |  |
| 1996-0213 | 0.66 | 0.74 | 0.78 |  |
| 0927-0159 | 0.67 | 0.76 | 0.8 |  |
| 3229-2671 | 0.67 | 0.79 | 0.81 |  |
| C276-0402 | 0.67 |  |  |  |
| CER0127724 | 0.67 |  |  |  |
| 3019-0037 | 0.68 |  |  |  |
| 4789-3825 | 0.68 |  |  |  |
| 1015-0074 | 0.68 |  |  |  |
| 0873-0090 | 0.68 |  |  |  |
| 3215-0601 | 0.68 |  |  |  |
| 2083-1978 | 0.68 | 0.77 |  |  |
| 1589-0140 | 0.68 | 0.72 | 0.8 | 0.81 |
| 0466-0074 | 0.68 | 0.73 | 0.73 | 0.8 |
| 2047-9619 | 0.69 |  |  |  |
| 0489-0004 | 0.69 |  |  |  |
| E470-0570 | 0.69 |  |  |  |
| 3046-9186 | 0.69 |  |  |  |
| 330004 | 0.69 | 0.63 |  |  |
| E013-0190 | 0.7 |  |  |  |
| K233-0118 | 0.7 |  |  |  |
| 1809-0088 | 0.71 |  |  |  |
| CER0126941 | 0.71 | 0.79 | 0.82 |  |
| E781-0440 | 0.71 |  |  |  |
| CER0128108 | 0.71 |  |  |  |
| 3202-0294 | 0.71 | 0.81 |  |  |
| CER0147308 | 0.72 |  |  |  |
| K086-0063 | 0.72 |  | 0.81 |  |
| 4817-0506 | 0.72 |  |  |  |
| 6233-0414 | 0.72 |  |  |  |
| CER0147232 | 0.73 |  | 0.82 |  |
| 4031-2881 | 0.73 |  |  |  |
| 1503938 | 0.73 |  |  |  |
| E208-0123 | 0.73 |  |  |  |
| 7100-1461 | 0.74 |  |  |  |
| 3229-2580 | 0.74 |  |  |  |
| 1885-0189 | 0.74 |  |  |  |
| CER0129015 | 0.74 |  |  |  |
| CER0114167 | 0.74 |  |  |  |
| 1500674 | 0.75 |  |  |  |
| 3454-2953 | 0.75 | 0.81 |  |  |
| 7218-1610 | 0.75 |  |  |  |
| 4031-2751 | 0.75 |  |  |  |
| 2806-0384 | 0.76 |  |  |  |
| 0873-0096 | 0.76 |  |  |  |
| 6239-0571 | 0.76 |  |  |  |
| CER0126949 | 0.77 |  |  |  |
| 7213-0566 | 0.77 |  |  |  |
| 8003-7417 | 0.77 |  |  |  |
| CER0128042 | 0.77 |  |  |  |
| CER0128841 | 0.77 |  |  |  |
| CER0129114 | 0.77 |  |  |  |
| 8003-0330 | 0.77 |  |  |  |
| 2437-0005 | 0.78 | 0.81 |  |  |
| K016-0004 | 0.79 |  |  |  |
| CER0129339 | 0.79 |  |  |  |
| 7011-1159 | 0.8 |  |  |  |
| 6228-1372 | 0.8 |  |  |  |
| CER0133694 | 0.8 |  |  |  |
| 8009-7508 | 0.8 |  |  |  |
| 1094-0243 | 0.81 |  | 0.81 |  |
| 1588-0183 | 0.81 | 0.74 | 0.81 |  |
| 8008-7266 | 0.81 |  |  |  |
| 2188-1693 | 0.81 |  |  |  |
| C794-0263 | 0.81 |  |  |  |
| E781-0038 | 0.81 |  |  |  |
| K890-0043 | 0.82 |  |  |  |
| 2039-0532 | 0.82 |  |  |  |
| CER0127673 | 0.82 |  |  |  |
| 1505138 | 0.82 | 0.76 | 0.83 | 0.82 |
| 1505298 | 0.82 |  |  |  |
| 5618-4188 |  | 0.61 | 0.76 | 0.8 |
| 8004-1113 |  | 0.62 | 0.79 | 0.68 |
| 8013-1492 |  | 0.71 | 0.81 | 0.82 |
| 8005-6924 |  |  | 0.78 | 0.78 |
| C243-0285 |  |  | 0.77 | 0.75 |
| 1505139 |  |  | 0.78 | 0.82 |
| 1503244 |  | 0.56 |  |  |
| 1503602 |  | 0.62 |  |  |
| 1500718 |  | 0.72 |  |  |
| 8007-7458 |  | 0.72 |  |  |
| 201650 |  | 0.72 |  |  |
| 8009-6343 |  | 0.75 |  |  |
| 1800031 |  | 0.77 |  |  |
| 2545-0104 |  | 0.78 |  |  |
| 1500328 |  | 0.79 |  |  |
| 0873-0036 |  | 0.79 |  |  |
| CER0128154 |  | 0.79 |  |  |
| 3388-0876 |  | 0.79 |  |  |
| 300554 |  | 0.81 |  |  |
| 8016-0477 |  | 0.82 |  |  |
| 0873-0098 |  | 0.82 |  |  |
| 1505149 |  |  | 0.73 |  |
| 6233-0162 |  |  | 0.78 |  |
| CER0131880 |  |  | 0.79 |  |
| 1996-0204 |  |  | 0.8 |  |
| 8010-2811 |  |  | 0.82 |  |
| 3172-0726 |  |  | 0.82 |  |
| 1488-0247 |  |  |  | 0.74 |
| C794-0039 |  |  |  | 0.81 |
| 8002-9170 |  |  |  | 0.82 |
| 1150-0014 |  |  |  | 0.82 |

**Figure 3.**
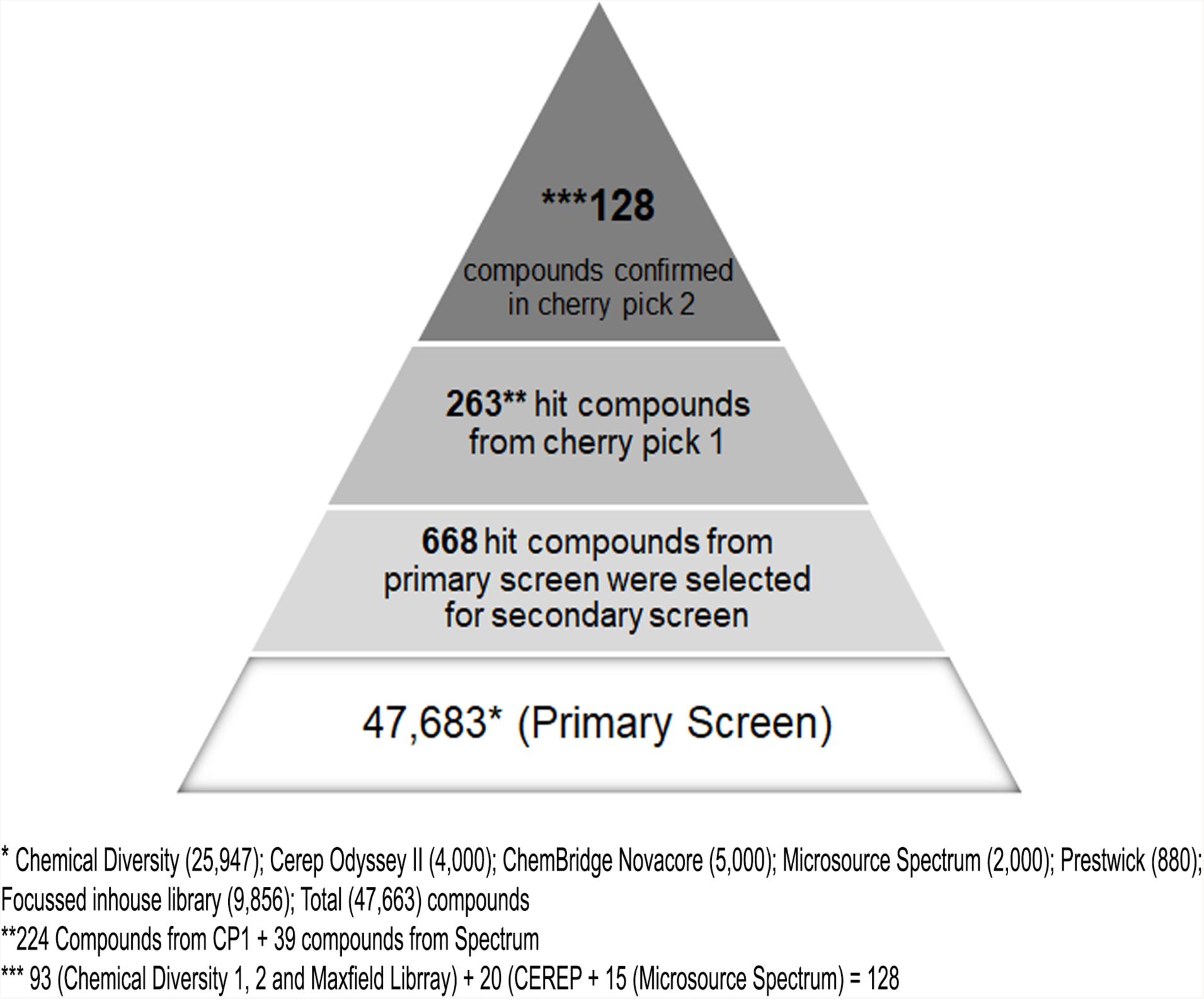
Schematic of screening results. Pyramid summarizing entire screen.

Shown in Figure 4A are the chemical structures of three compounds effective at all four doses tested. Table 4 shows the structures of 128 hit compounds with is formula and molecular weight.

**Table 4.**
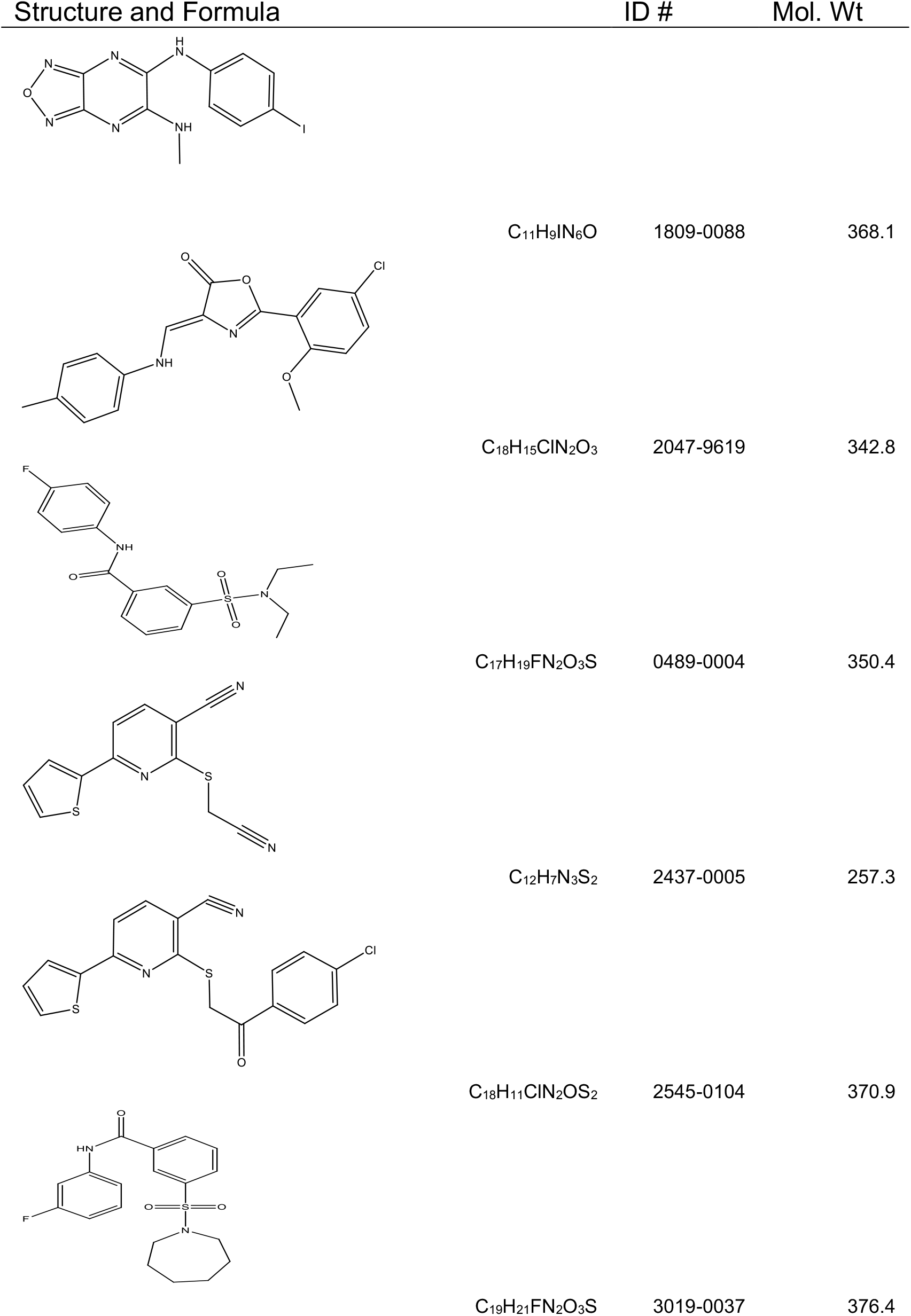
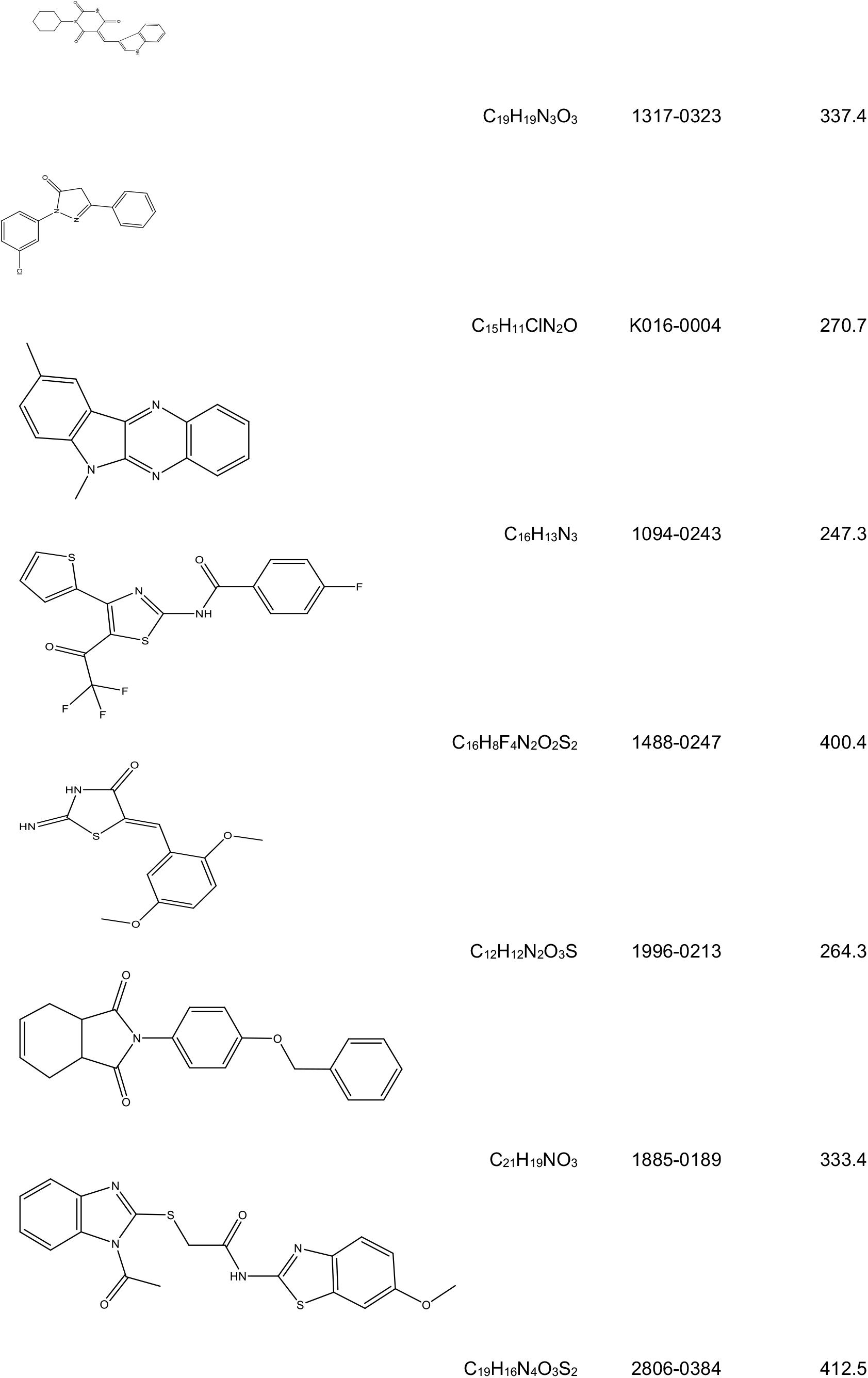
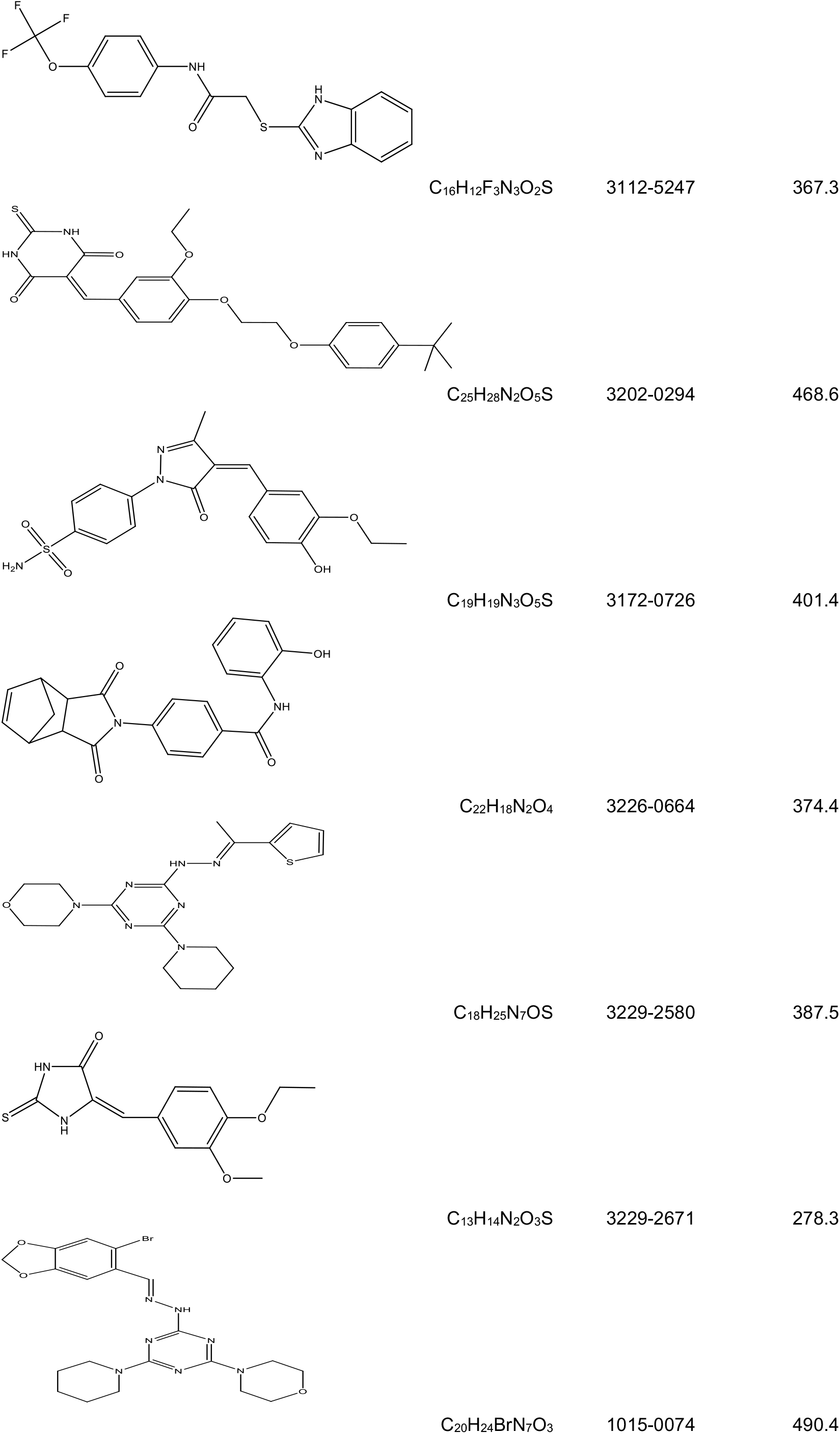
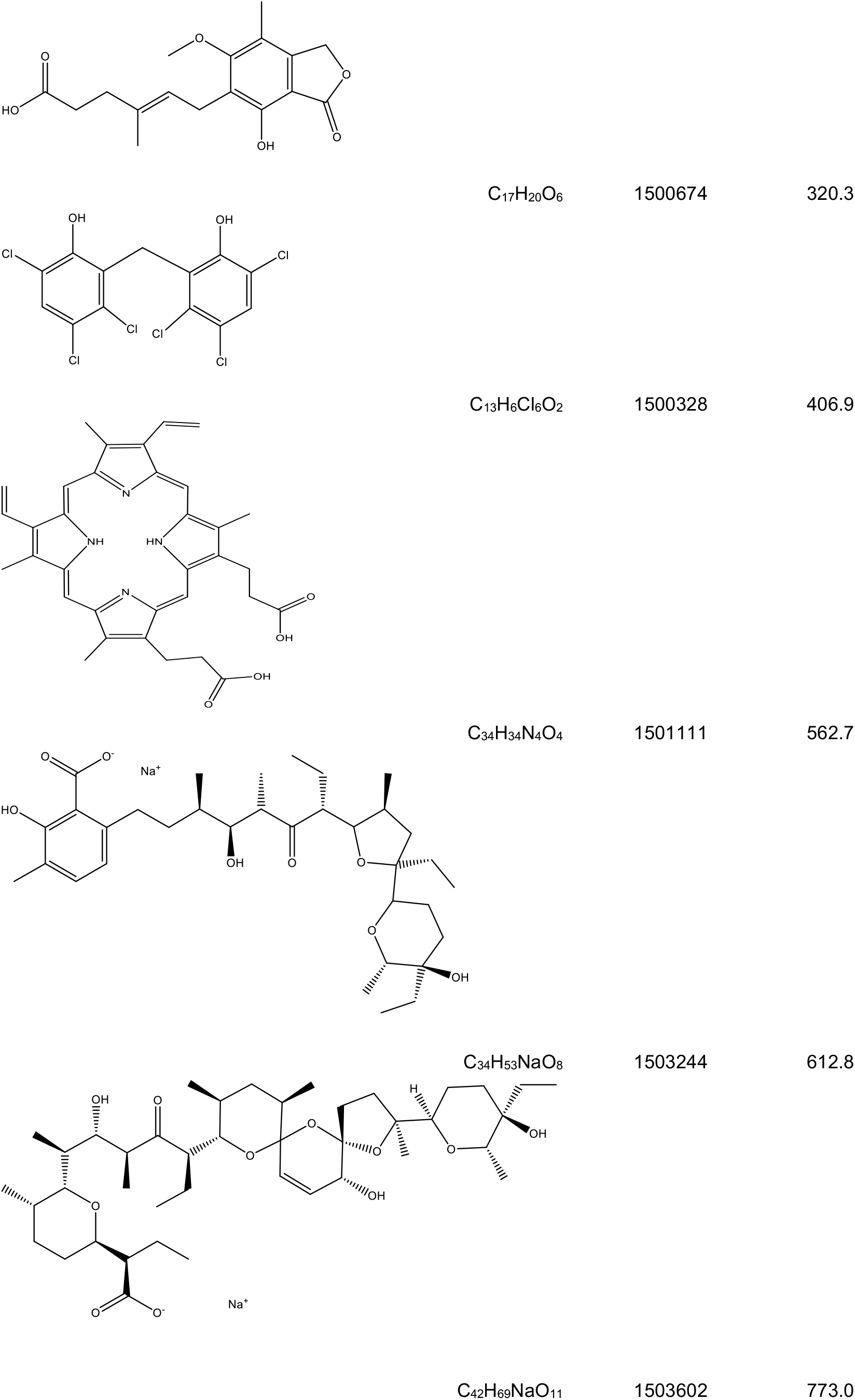
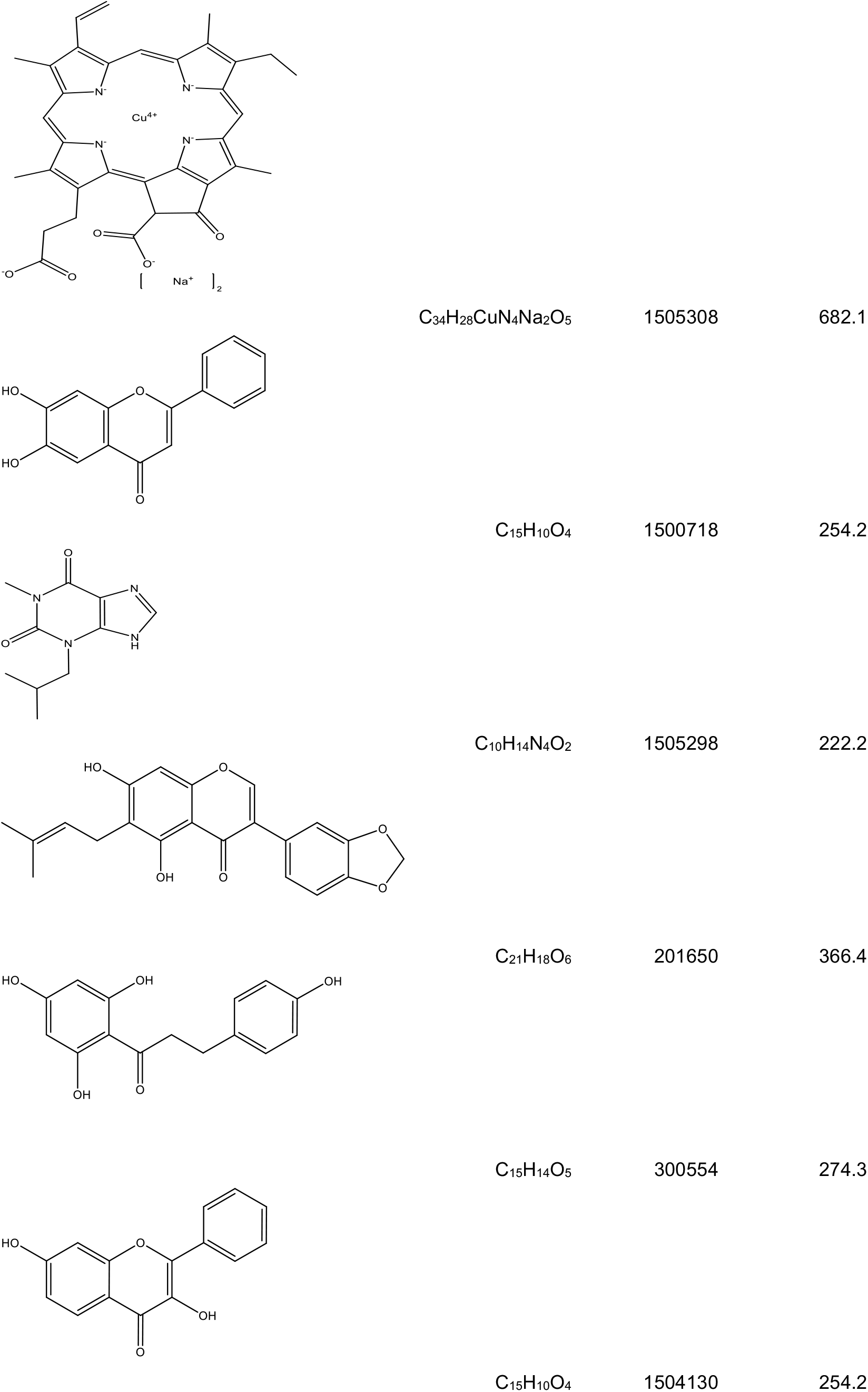
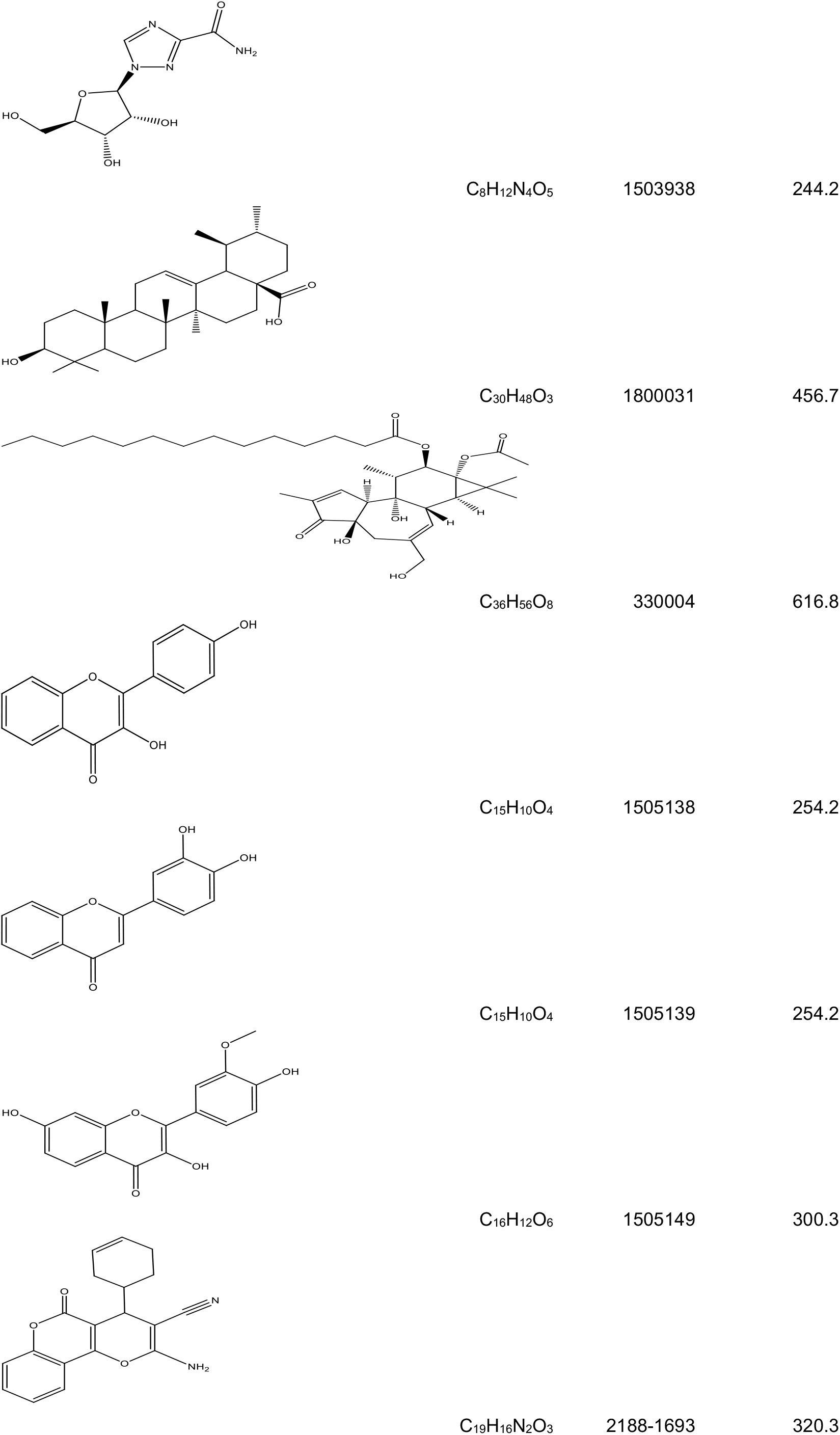
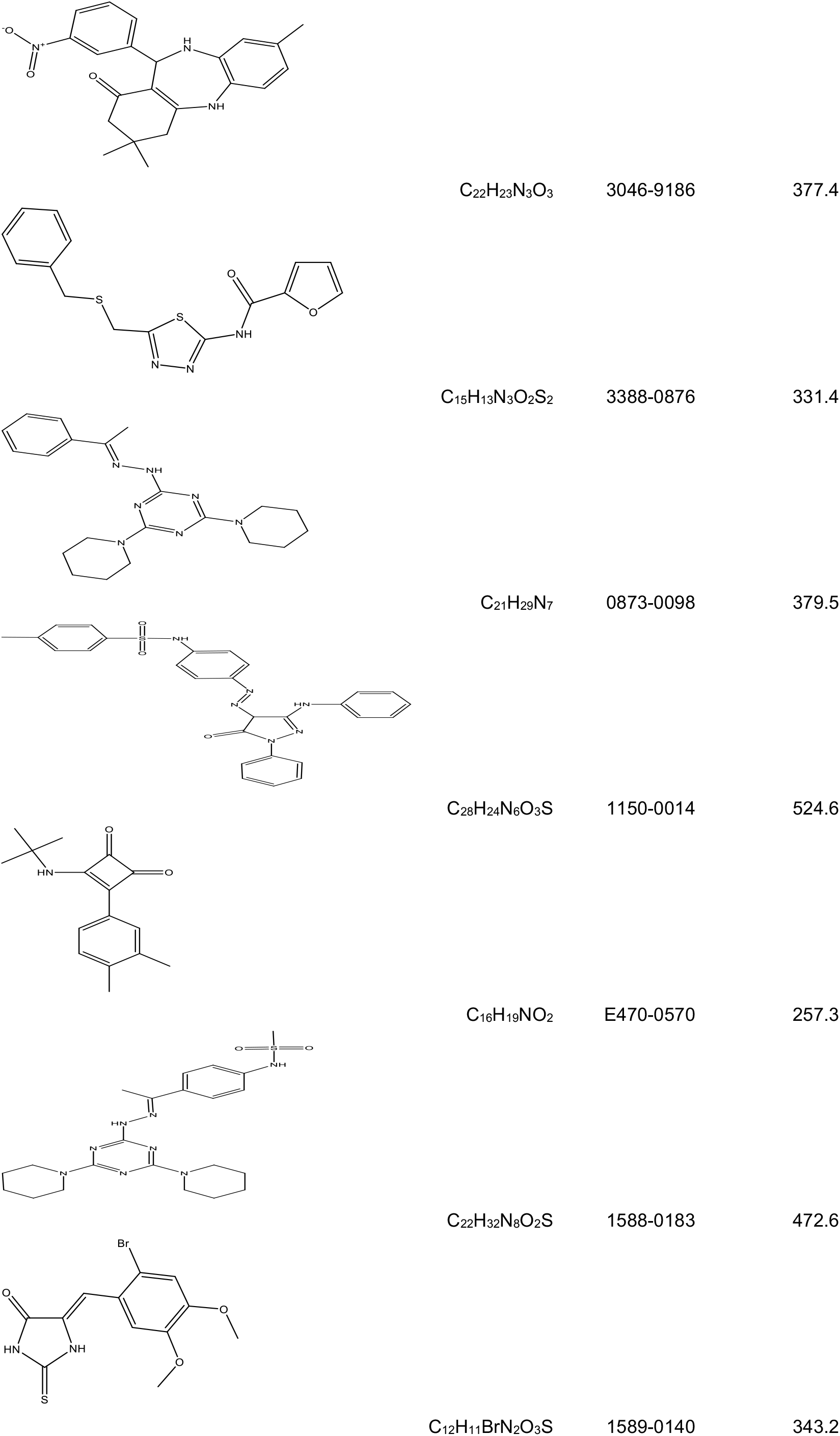
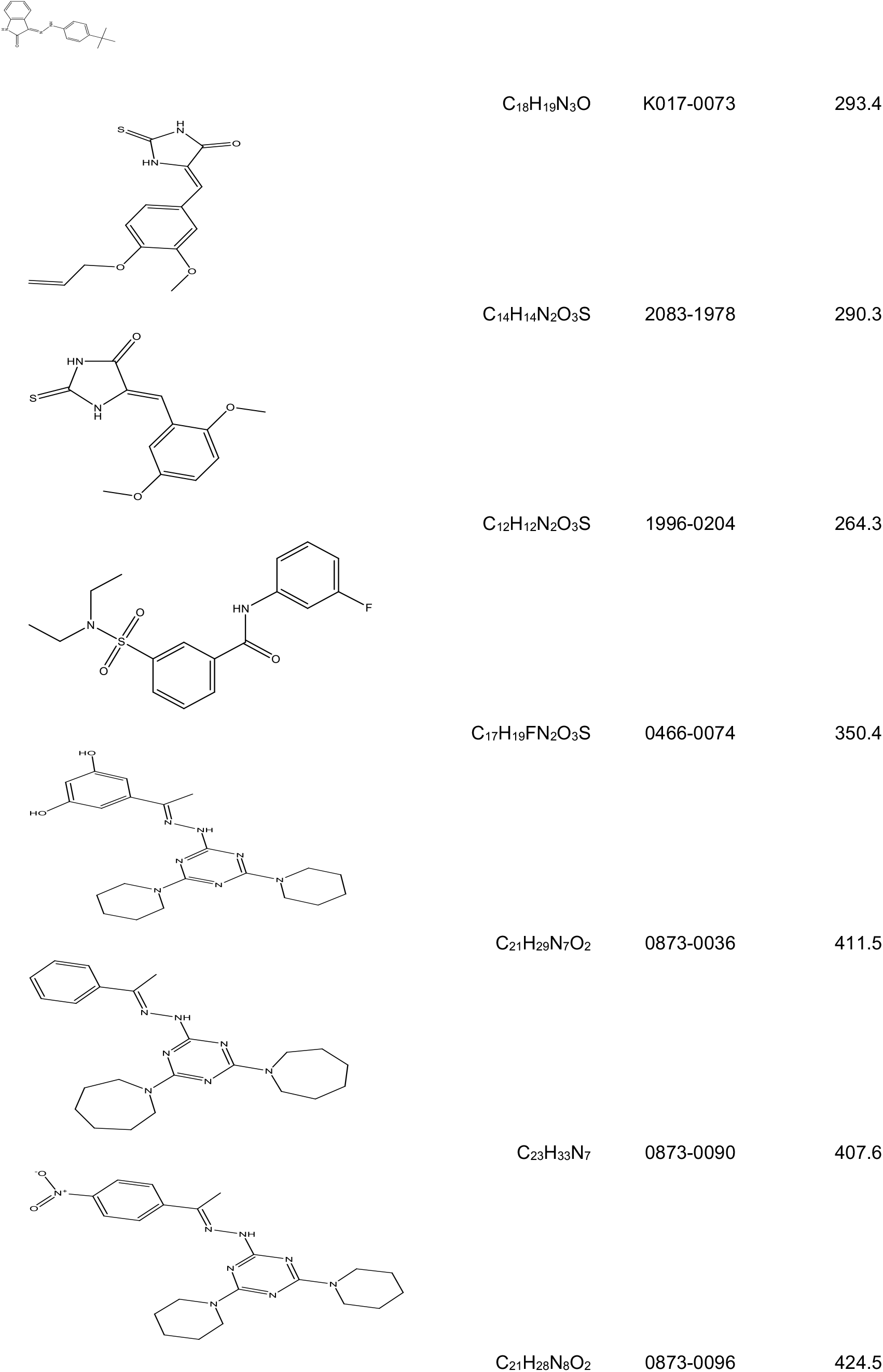
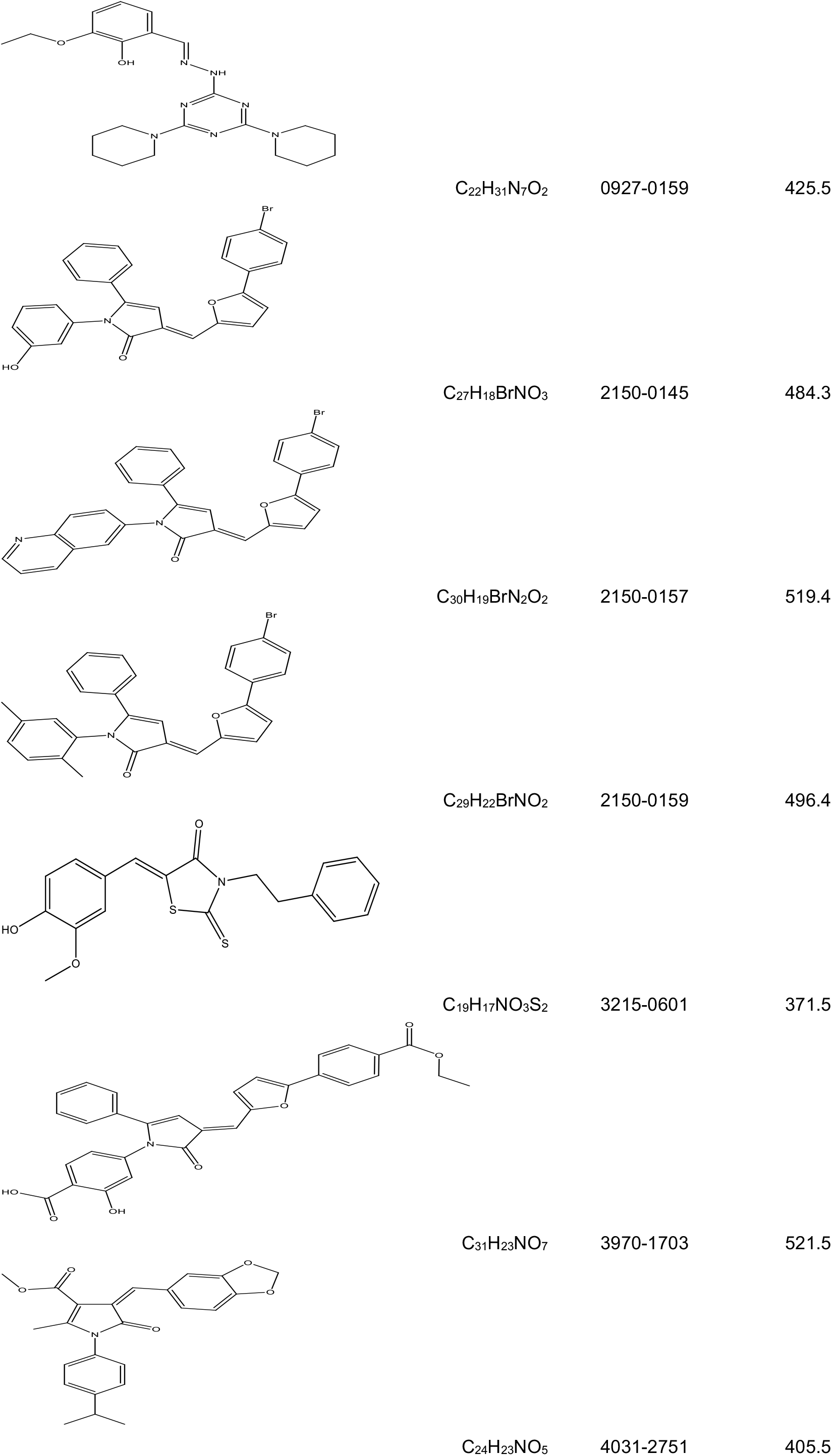
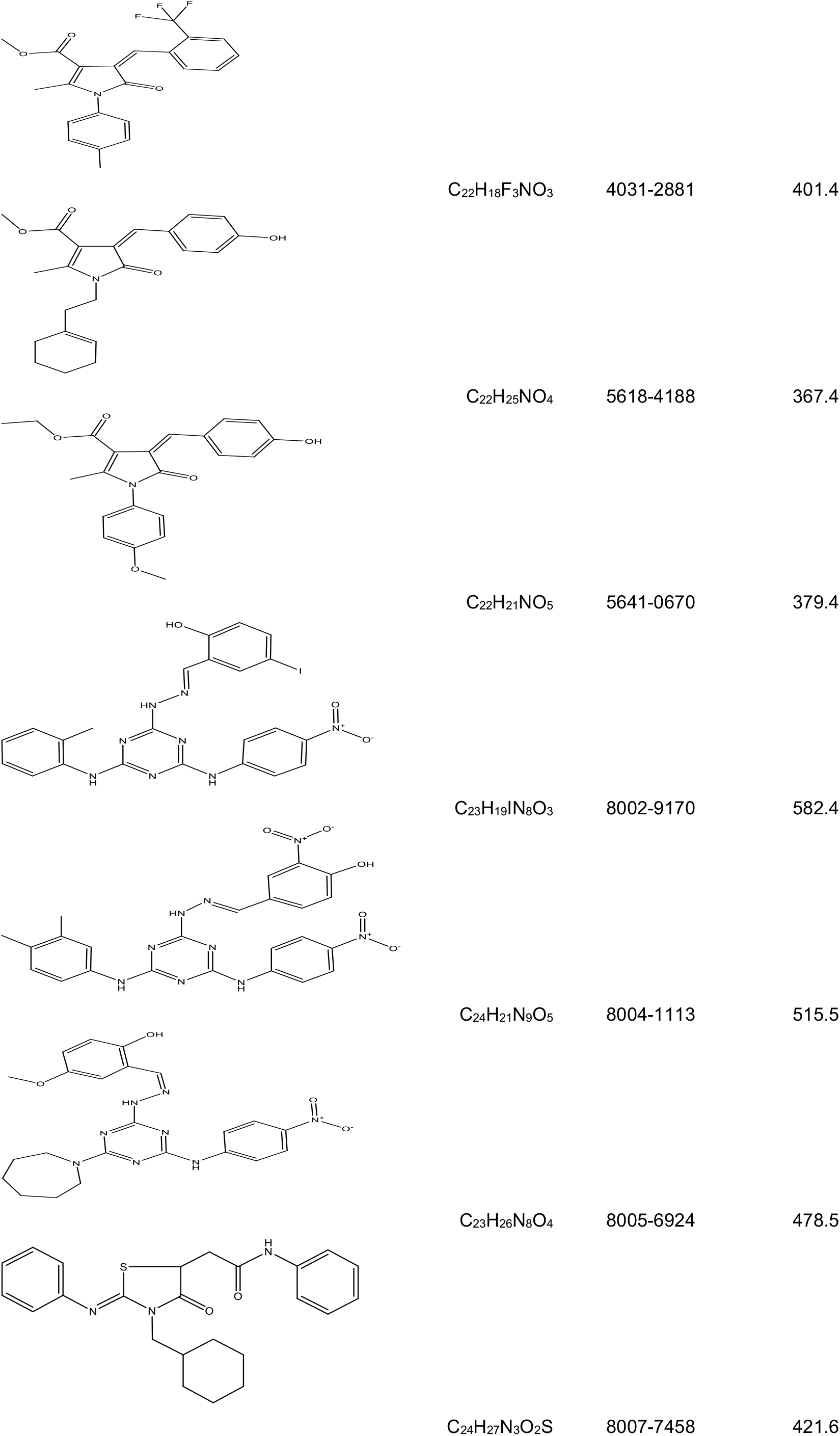
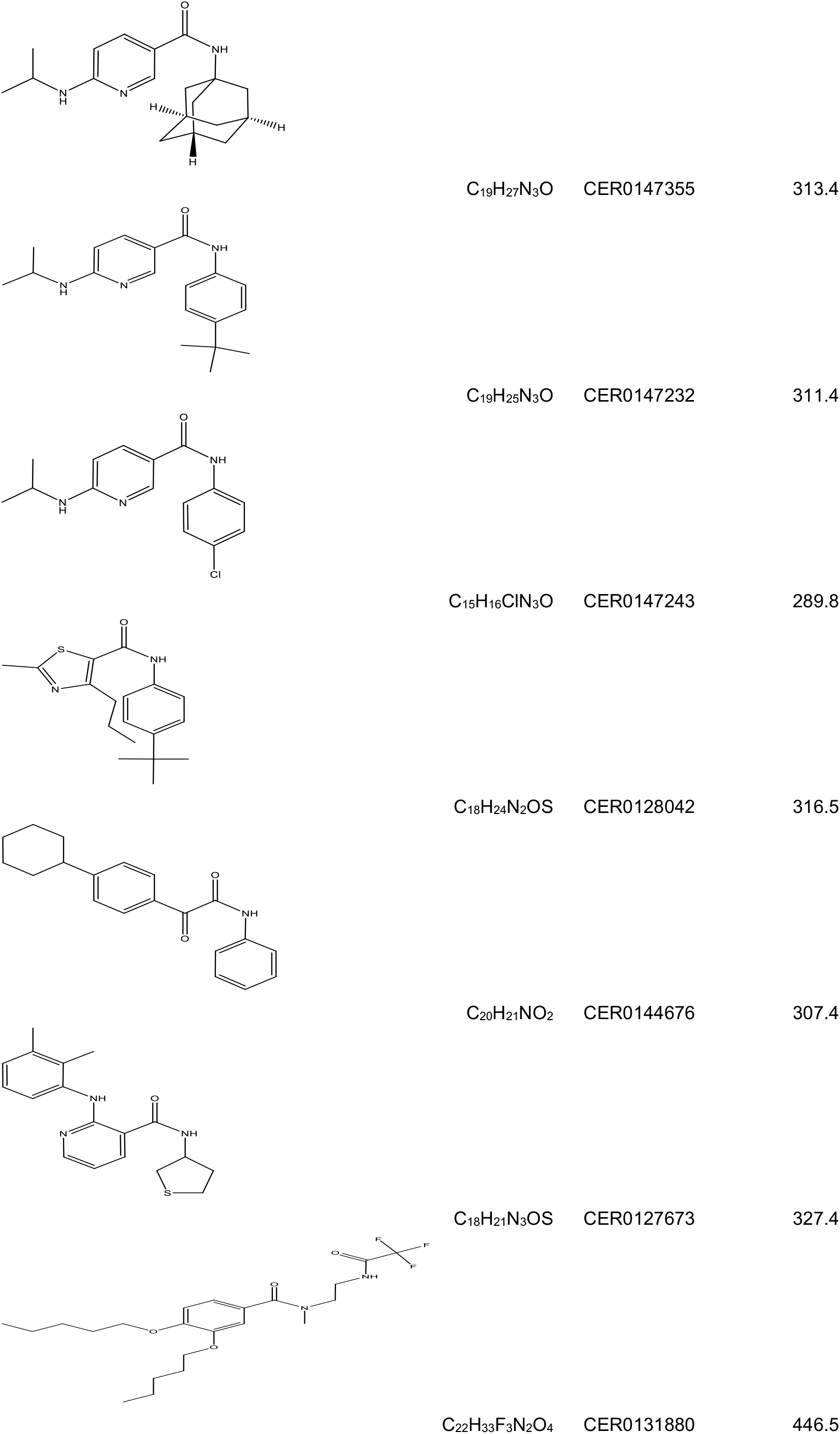
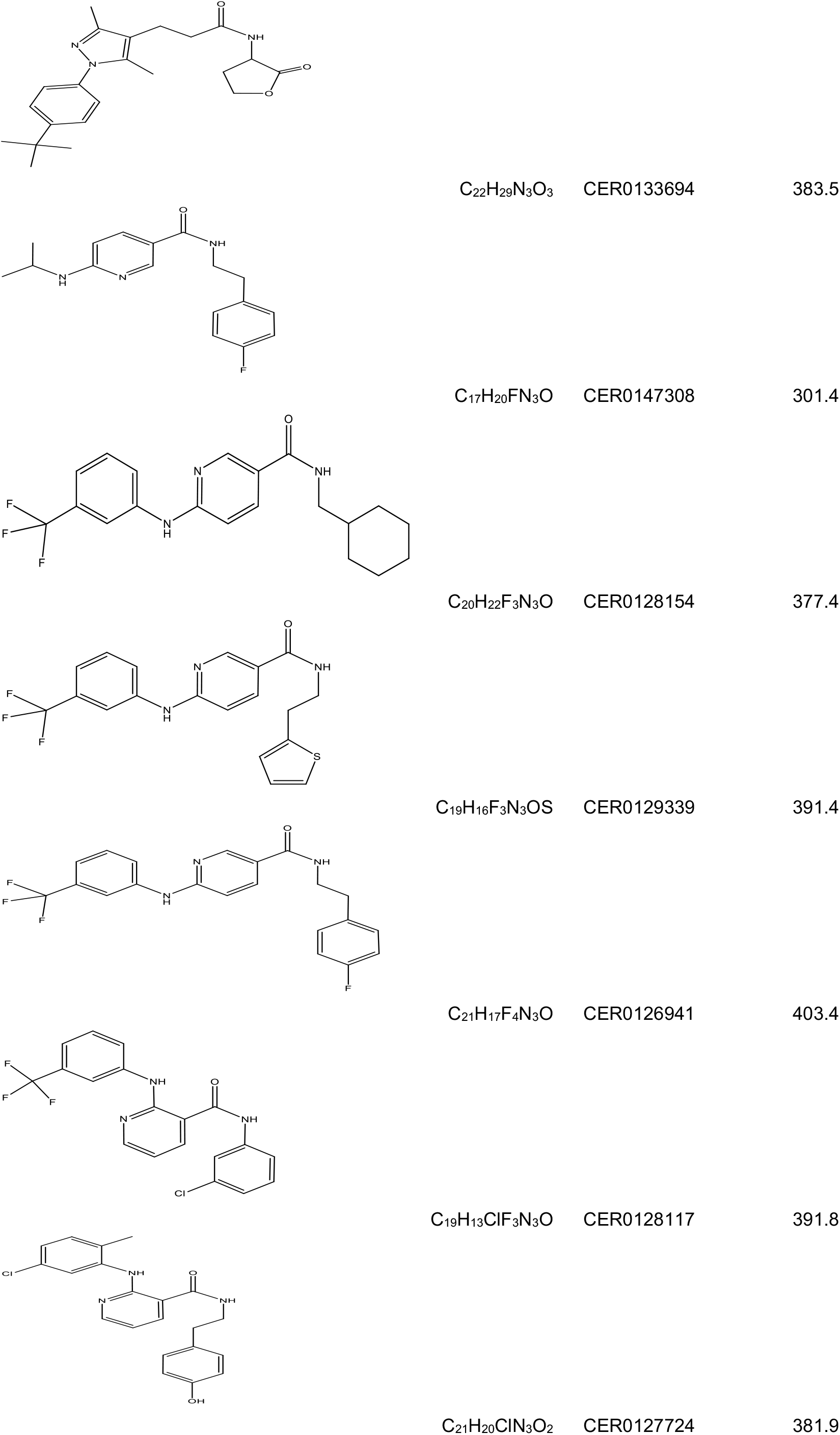
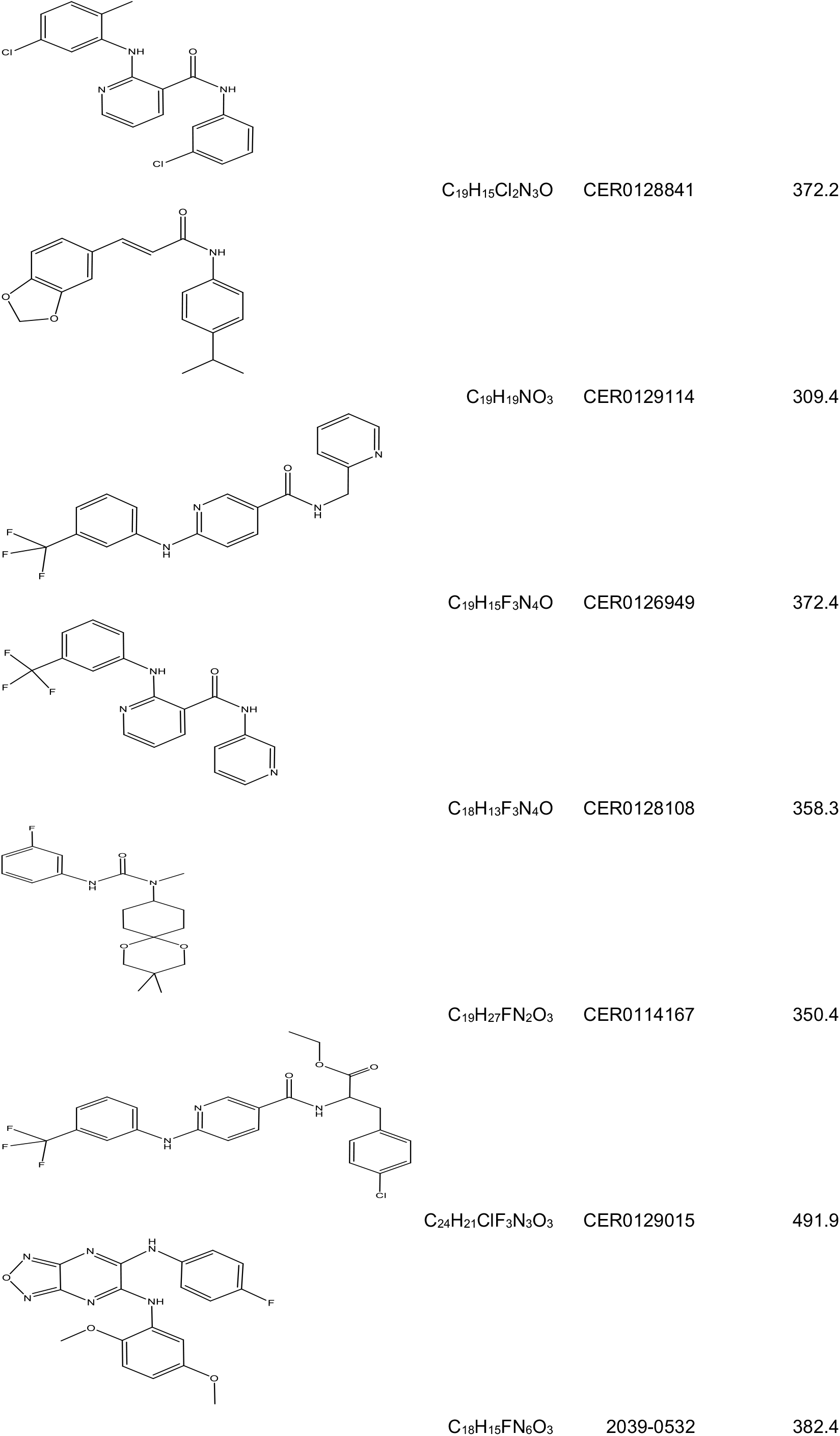
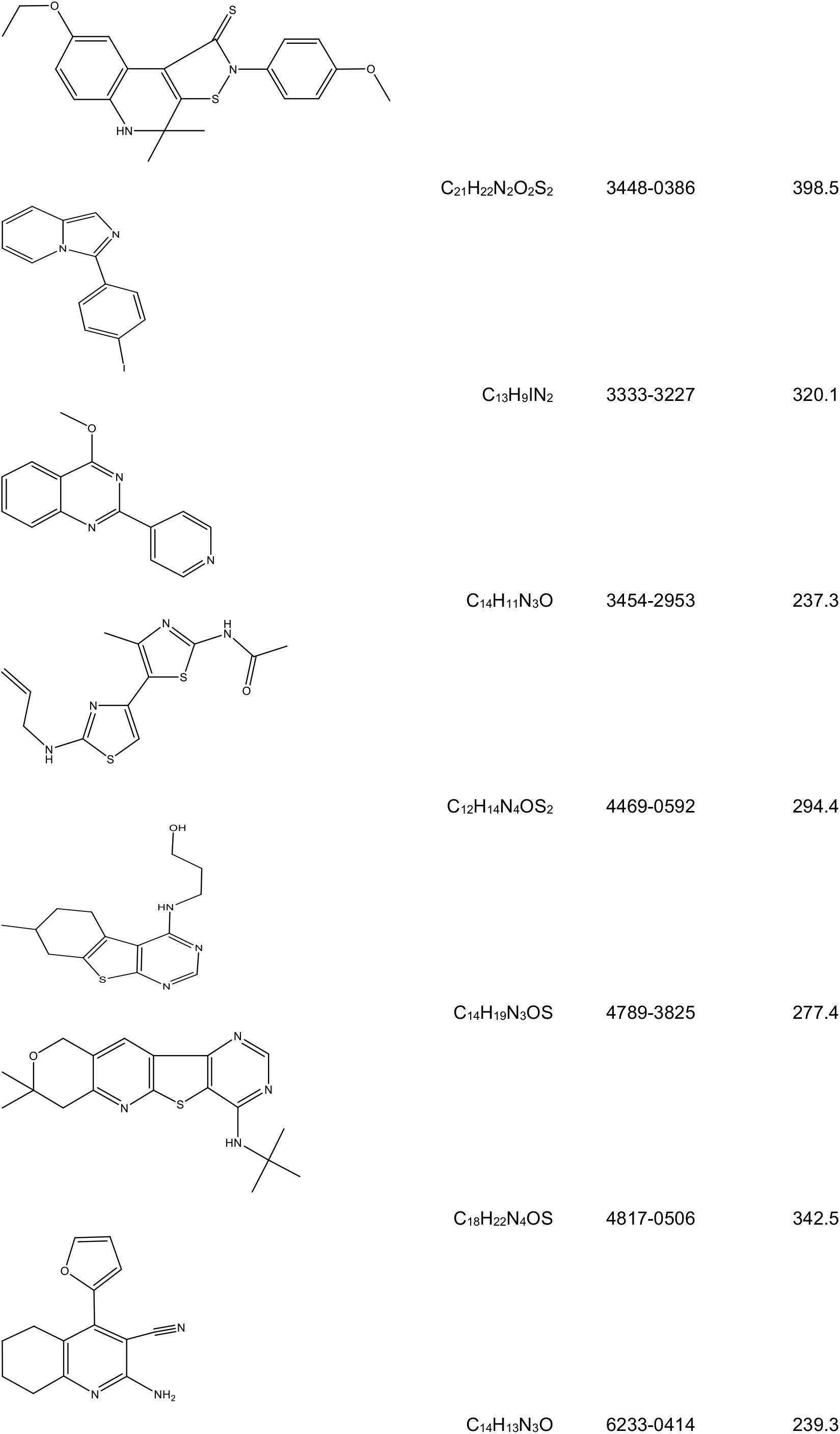
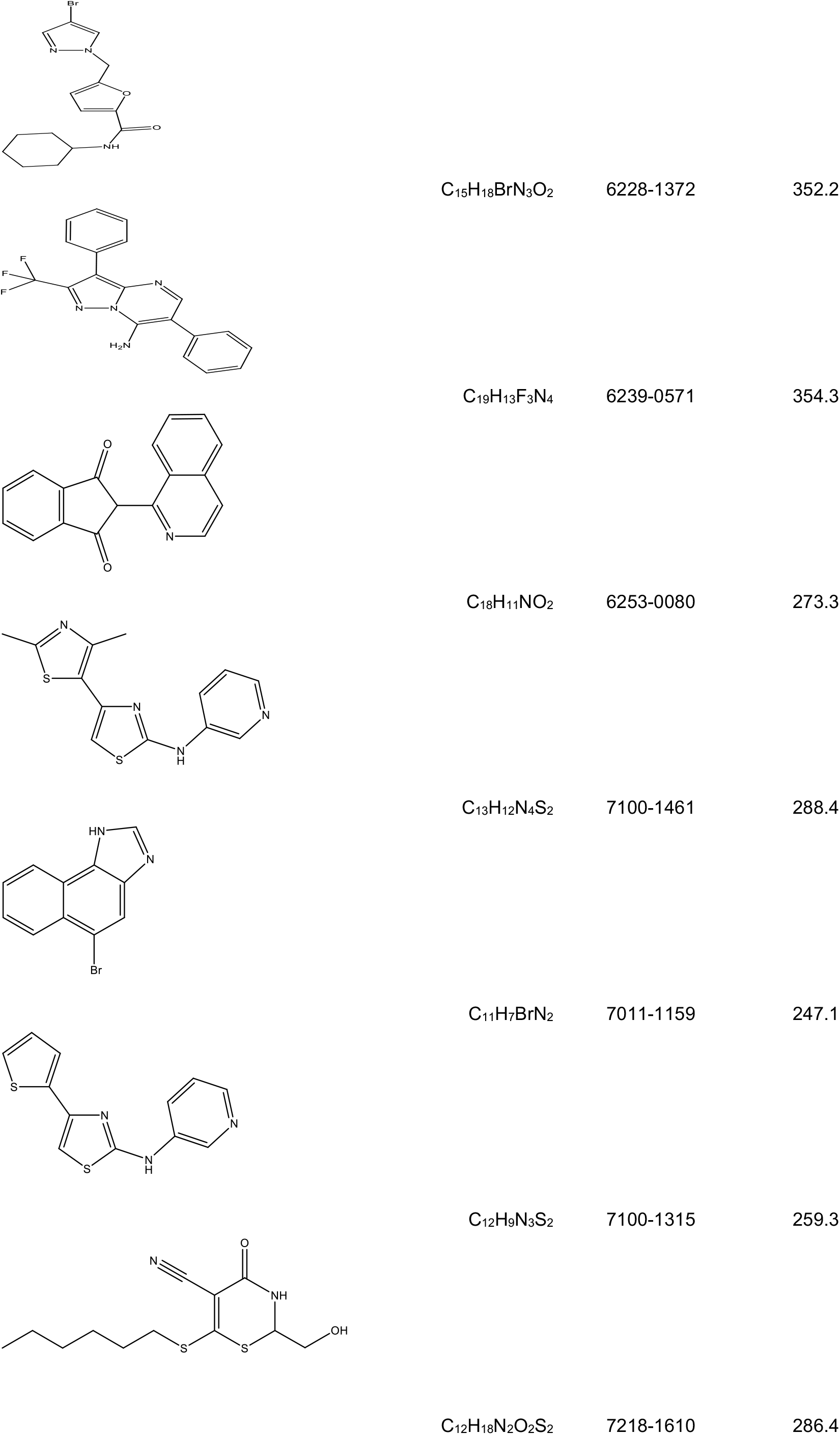
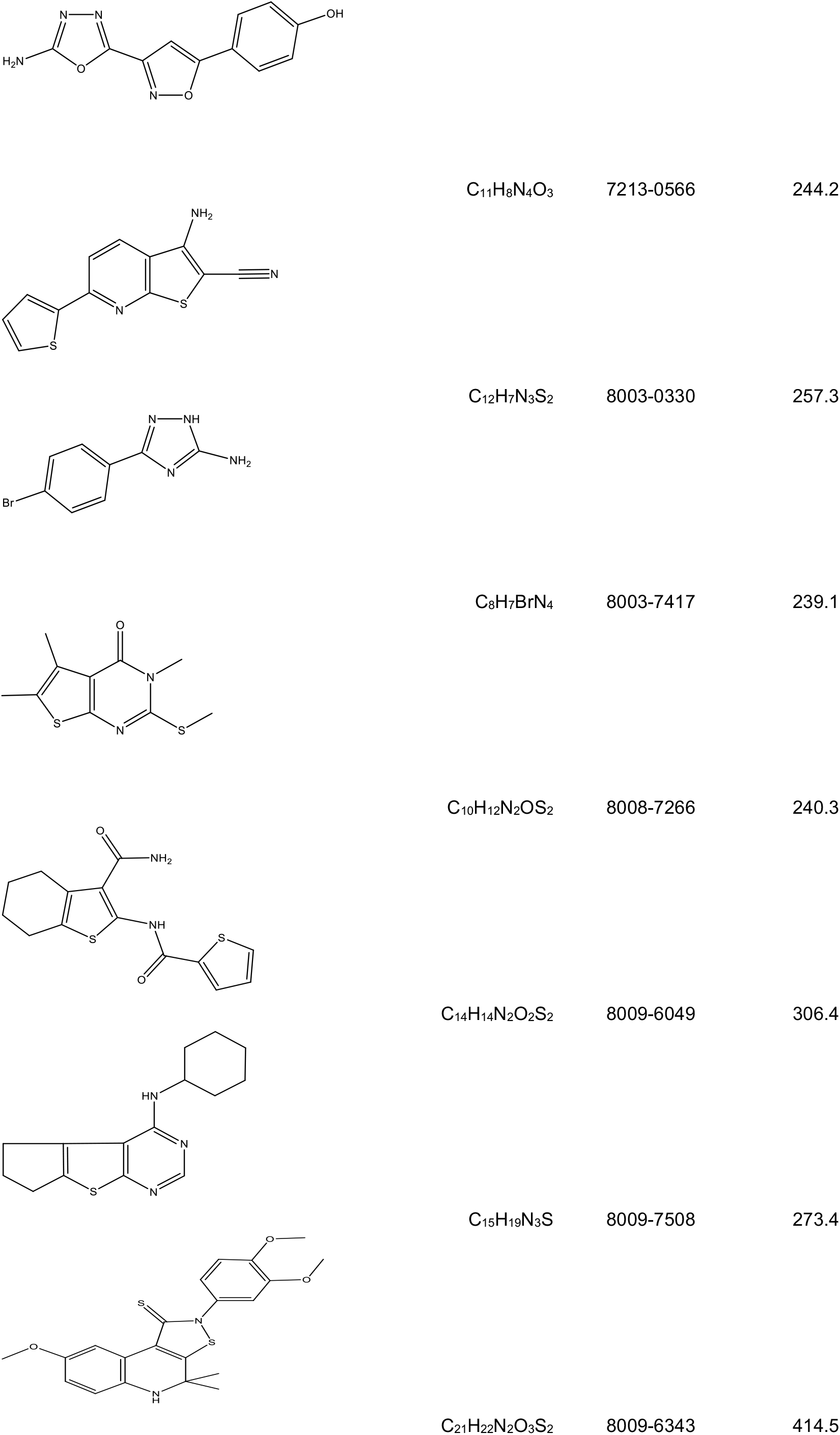
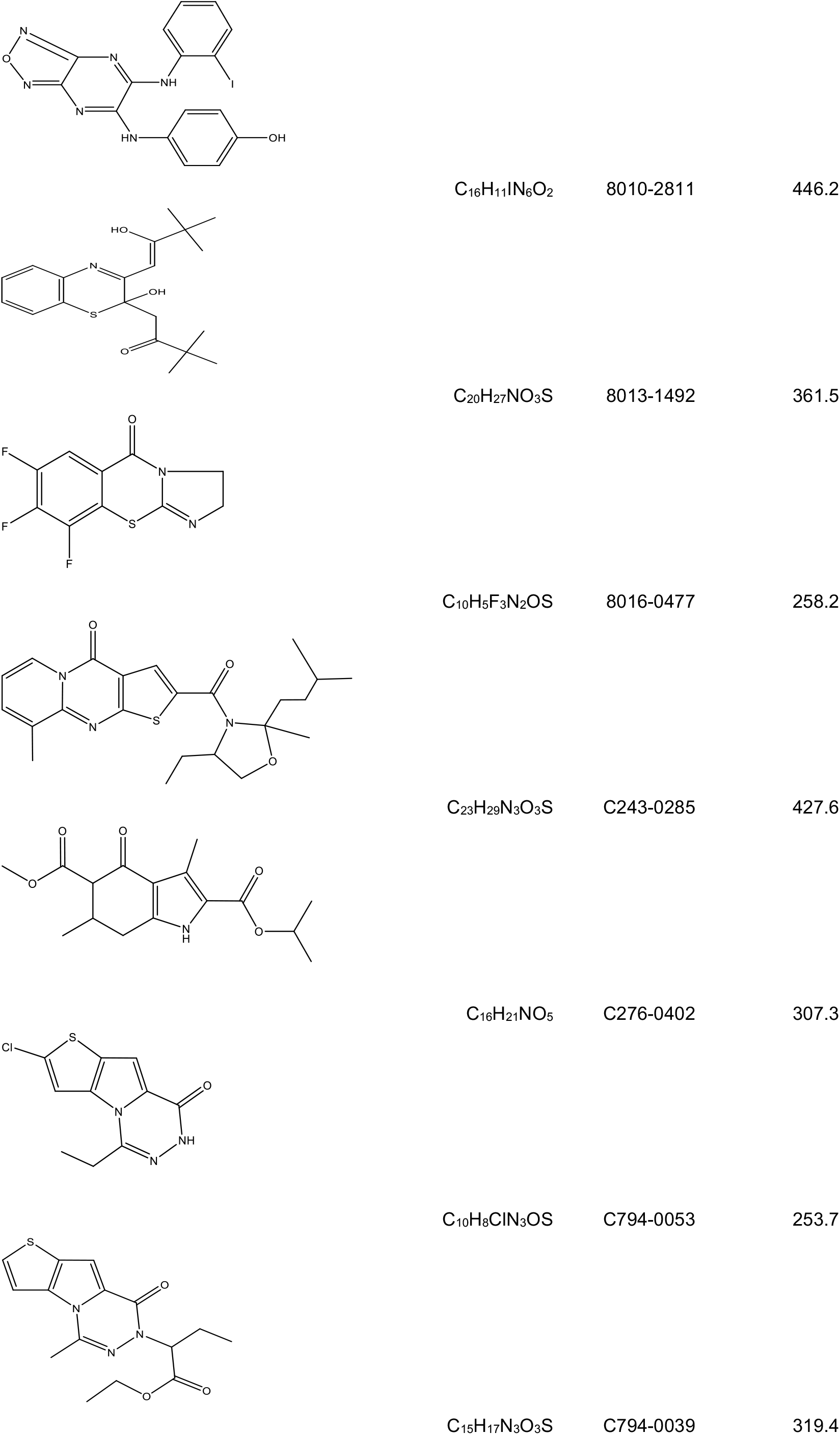
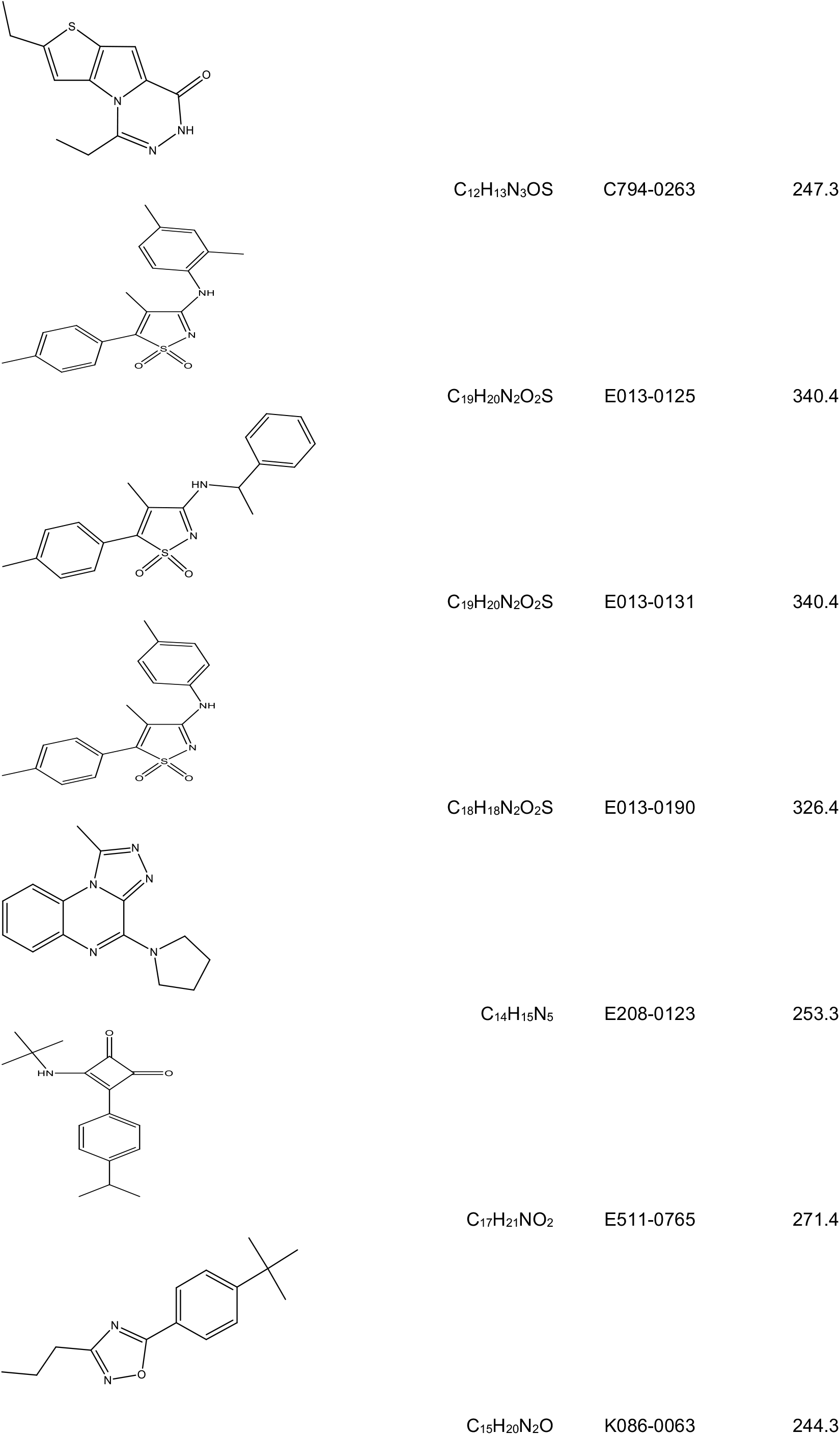
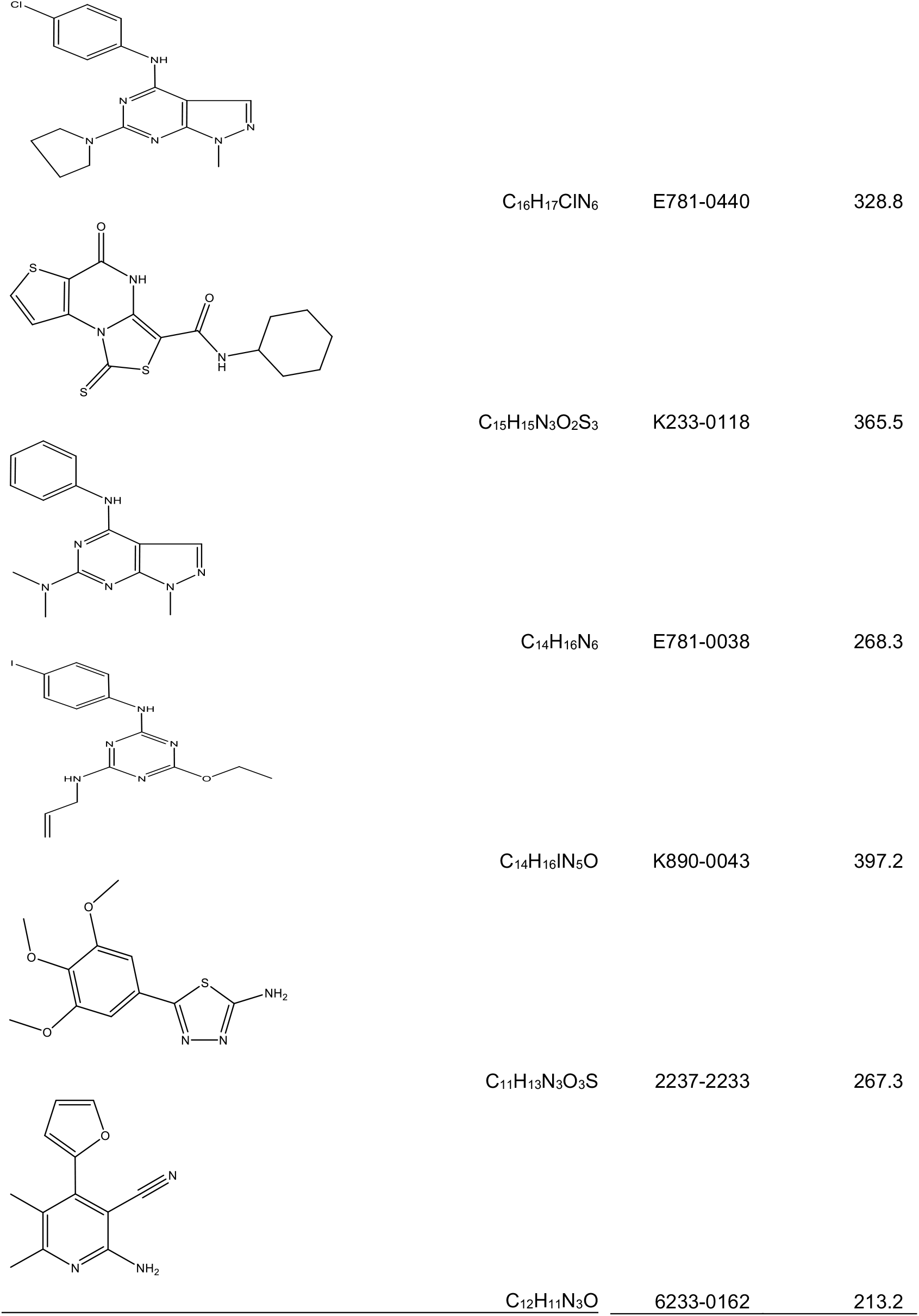
Structures of all hit compounds.

**Figure 4.**
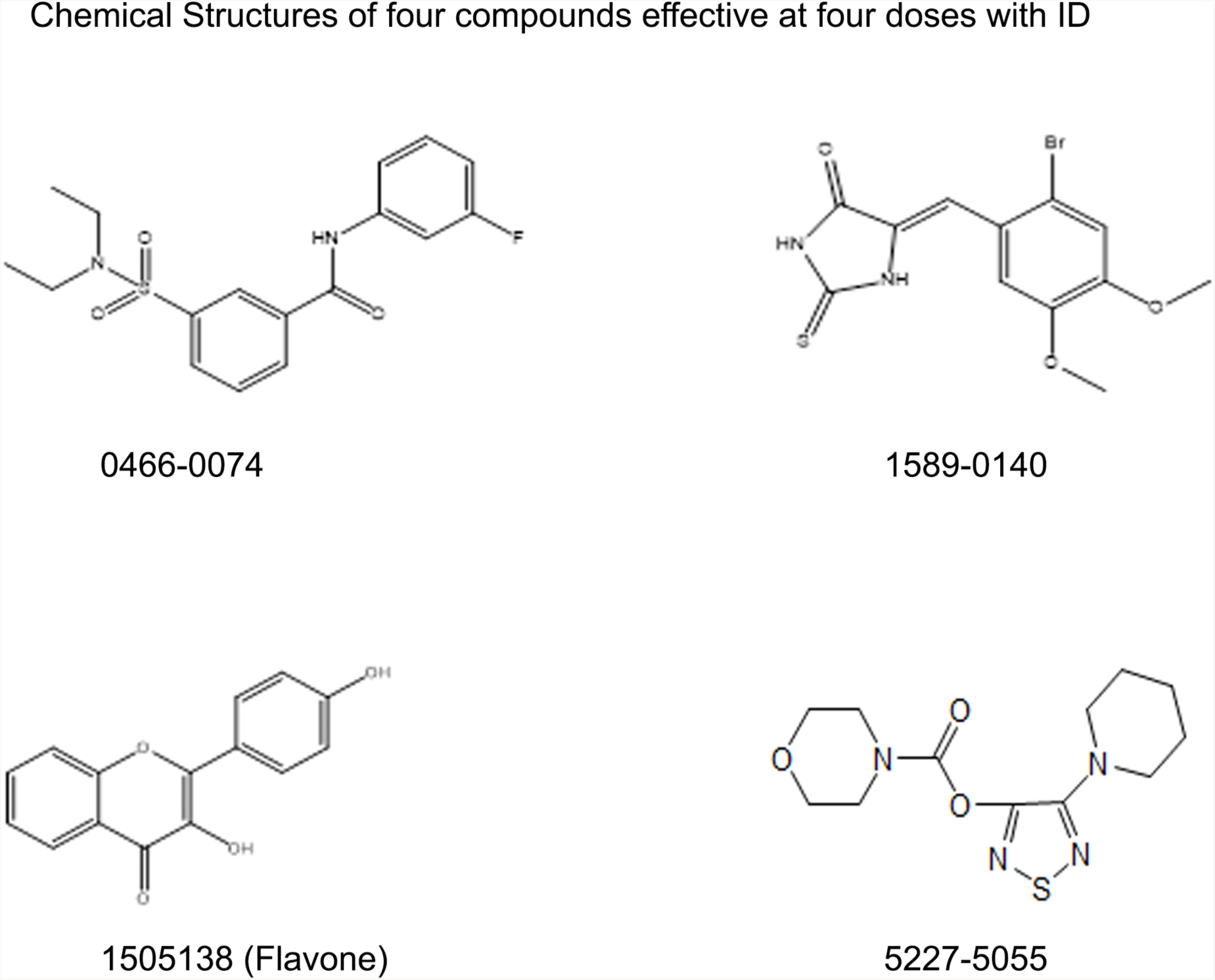
Chemical structures of three compounds effective at all four doses tested.

## Discussion

We screened NPC1 mutant primary fibroblasts bearing a heterozygous I1061T mutation using a previously established automated screening protocol to identify compounds that partially reverse the NPC1 phenotype (24). We exploited the selective cholesterol binding property of filipin for the fluorescence microscopy-based assay. The classic characteristic of NPC1 mutant cells unlike WT cells is the accumulation of free cholesterol in LSOs, organelles that are related to late endosomes but also contain protein markers that are usually not abundant in late endosomes (30). The mutant form of NPC proteins (NPC1 and NPC2) causes a defect in efflux of cholesterol from late endosomes, resulting in high levels of cholesterol in the LSOs.

Using our previously described methods (24), we identified 668 compounds from a diverse library of 47,683 compounds from six different vendors after primary screening. These compounds significantly decreased the filipin staining in the compartment at 10 µM and were not cytotoxic. The 668 hits from primary screen were cherry picked in a secondary screen, which resulted in confirming 263 compounds. These 263 confirmed hits were further analyzed in a second round and tested at 4 different doses (10, 1, 0.1 and 0.03 µM). The second round of analysis reconfirmed 134 compounds that corrected the NPC1 phenotype, as seen by reduced LSO filipin signal at various concentrations. Some of the strong hit compounds were rejected based on information available in literature and its relevance to the goal of this screen. For instance, monensin, which increases lysosomal pH (31, 32) and is toxic was rejected. It has also been reported that monensin interrupts recycling of LDL receptors (33). Another compound that was disqualified was Protoporphyrin IX (PPIX) – a heme transporter that completely abrogated cholesterol accumulation in the NPC1 fibroblasts. The activity of PPIX in reducing cholesterol accumulation can be explained, based on previous finding that PPIX disrupts phospholipid, bile acid and cholesterol equilibrium in hepatocytes and Kupffer cells (34). Since PPIX perturbs heme transport, it will have an adverse effect on several other biological processes and is not a good drug candidate.

One of the hits, Compound 2-a-1, was effective at three concentrations (10, 3.33 and 1.11 uM). We have previously characterized this compound on CT60 cells, and we found it reduces the uptake of DiI-LDL, indicating that it might be down regulating LDL receptors (29, 35) and cholesterol accumulation. Inhibition of cholesterol uptake leads to exclusion of this drug.

Our library contained combinatorially synthesized small molecules, FDA approved compounds, and natural product compounds. We observed several hits from the natural product library, which had been previously reported to have cholesterol reducing activity. For instance, one of the hit compounds is a plant triterpenoid ursolic acid, which has been shown to be a PPAR-agonist and significantly reduces intra-cellular cholesterol in hepatocytes (36). Another class of hit compounds were flavones, belonging to a family of flavanoids- a diverse polyphenolic compounds found in the plant kingdom (37). In the past, flavones have been suggested to have anti-athersclerotic and cholesterol lowering activity in several studies. However, like HDACi’s, flavonoids affect multiple genes in the cells resulting in many off-target effects, making them unsuitable as a drug candidate.

In conclusion, the studies presented here show that small molecules added to cells can partially correct the NPC1 defect in cultured cells. Since several of these compounds are effective in reducing cholesterol accumulation at concentrations at which they are non-toxic to cultured NPC1 cells, they have potential use as lead compounds for drug discovery.

## Acknowledgements

This research was funded by grants from the Ara Parseghian Medical Research Foundation.

## References

1. Carstea ED, Morris JA, Coleman KG, Loftus SK, Zhang D, Cummings C, et al. Niemann-Pick C1 disease gene: homology to mediators of cholesterol homeostasis. Science. 1997;277(5323):228–31.

2. Sun X, Marks DL, Park WD, Wheatley CL, Puri V, O’Brien JF, et al. Niemann-Pick C variant detection by altered sphingolipid trafficking and correlation with mutations within a specific domain of NPC1. Am J Hum Genet. 2001;68(6):1361–72.

3. Mukherjee S, Maxfield FR. Lipid and cholesterol trafficking in NPC. Biochim Biophys Acta. 2004;1685(1-3):28–37.

4. Fink JK, Filling-Katz MR, Sokol J, Cogan DG, Pikus A, Sonies B, et al. Clinical spectrum of Niemann-Pick disease type C. Neurology. 1989;39(8):1040–9.

5. Sturley SL, Patterson MC, Balch W, Liscum L. The pathophysiology and mechanisms of NP-C disease. Biochim Biophys Acta. 2004;1685(1-3):83–7.

6. Greenberg CR, Barnes JG, Kogan S, Seargeant LE. A rare case of Niemann-Pick disease type C without neurological involvement in a 66-year-old patient. Mol Genet Metab Rep. 2015;3:18–20.

7. Brown MS, Goldstein JL. A receptor-mediated pathway for cholesterol homeostasis. Science. 1986;232(4746):34–47.

8. Naureckiene S, Sleat DE, Lackland H, Fensom A, Vanier MT, Wattiaux R, et al. Identification of HE1 as the second gene of Niemann-Pick C disease. Science. 2000;290(5500):2298–301.

9. Watari H, Blanchette-Mackie EJ, Dwyer NK, Glick JM, Patel S, Neufeld EB, et al. Niemann-Pick C1 protein: obligatory roles for N-terminal domains and lysosomal targeting in cholesterol mobilization. Proc Natl Acad Sci U S A. 1999;96(3):805–10.

10. Qian H, Wu X, Du X, Yao X, Zhao X, Lee J, et al. Structural Basis of Low-pH-Dependent Lysosomal Cholesterol Egress by NPC1 and NPC2. Cell. 2020;182(1):98-111.e18.

11. Long T, Qi X, Hassan A, Liang Q, De Brabander JK, Li X. Structural basis for itraconazole-mediated NPC1 inhibition. Nat Commun. 2020;11(1):152.

12. Horenkamp FA, Valverde DP, Nunnari J, Reinisch KM. Molecular basis for sterol transport by StART-like lipid transfer domains. EMBO J. 2018;37(6).

13. Lim CY, Davis OB, Shin HR, Zhang J, Berdan CA, Jiang X, et al. ER-lysosome contacts enable cholesterol sensing by mTORC1 and drive aberrant growth signalling in Niemann-Pick type C. Nat Cell Biol. 2019;21(10):1206–18.

14. Meng Y, Heybrock S, Neculai D, Saftig P. Cholesterol Handling in Lysosomes and Beyond. Trends Cell Biol. 2020;30(6):452–66.

15. Mesmin B, Pipalia NH, Lund FW, Ramlall TF, Sokolov A, Eliezer D, et al. STARD4 abundance regulates sterol transport and sensing. Mol Biol Cell. 2011;22(21):4004–15.

16. Rosenbaum AI, Zhang G, Warren JD, Maxfield FR. Endocytosis of beta-cyclodextrins is responsible for cholesterol reduction in Niemann-Pick type C mutant cells. Proc Natl Acad Sci U S A. 107(12):5477–82.

17. Abi-Mosleh L, Infante RE, Radhakrishnan A, Goldstein JL, Brown MS. Cyclodextrin overcomes deficient lysosome-to-endoplasmic reticulum transport of cholesterol in Niemann-Pick type C cells. Proc Natl Acad Sci U S A. 2009;106(46):19316–21.

18. Liu B, Turley SD, Burns DK, Miller AM, Repa JJ, Dietschy JM. Reversal of defective lysosomal transport in NPC disease ameliorates liver dysfunction and neurodegeneration in the npc1-/-mouse. Proc Natl Acad Sci U S A. 2009;106(7):2377–82.

19. Matsuo M, Shraishi K, Wada K, Ishitsuka Y, Doi H, Maeda M, et al. Effects of intracerebroventricular administration of 2-hydroxypropyl-beta-cyclodextrin in a patient with Niemann-Pick Type C disease. Mol Genet Metab Rep. 2014;1:391–400.

20. Matsuo M, Togawa M, Hirabaru K, Mochinaga S, Narita A, Adachi M, et al. Effects of cyclodextrin in two patients with Niemann-Pick Type C disease. Mol Genet Metab. 2013;108(1):76–81.

21. Pipalia NH, Cosner CC, Huang A, Chatterjee A, Bourbon P, Farley N, et al. Histone deacetylase inhibitor treatment dramatically reduces cholesterol accumulation in Niemann-Pick type C1 mutant human fibroblasts. Proc Natl Acad Sci U S A. 2011;108(14):5620–5.

22. Munkacsi AB, Hammond N, Schneider RT, Senanayake DS, Higaki K, Lagutin K, et al. Normalization of Hepatic Homeostasis in the Npc1(nmf164) Mouse Model of Niemann-Pick Type C Disease Treated with the Histone Deacetylase Inhibitor Vorinostat. J Biol Chem. 2017;292(11):4395–410.

23. Pipalia NH, Subramanian K, Mao S, Ralph H, Hutt DM, Scott SM, et al. Histone deacetylase inhibitors correct the cholesterol storage defect in most Niemann-Pick C1 mutant cells. J Lipid Res. 2017;58(4):695–708.

24. Pipalia NH, Huang A, Ralph H, Rujoi M, Maxfield FR. Automated microscopy screening for compounds that partially revert cholesterol accumulation in Niemann-Pick C cells. J Lipid Res. 2006;47(2):284–301.

25. Zhang JH, Chung TD, Oldenburg KR. A Simple Statistical Parameter for Use in Evaluation and Validation of High Throughput Screening Assays. J Biomol Screen. 1999;4(2):67–73.

26. Patrick AD, Lake BD. Deficiency of an acid lipase in Wolman’s disease. Nature. 1969;222(5198):1067–8.

27. Rosenbaum AI, Rujoi M, Huang AY, D. H, Grabowski GA, Maxfield FR. Chemical screen to reduce sterol accumulation in Niemann-Pick C disease cells identifies novel lysosomal acid lipase inhibitors. Biochim Biophys Acta. 2009;1791(12):1155–65.

28. Patel JZ, Nevalainen TJ, Savinainen JR, Adams Y, Laitinen T, Runyon RS, et al. Optimization of 1,2,5-thiadiazole carbamates as potent and selective ABHD6 inhibitors. ChemMedChem. 2015;10(2):253–65.

29. Rujoi M, Pipalia NH, Maxfield FR. Cholesterol pathways affected by small molecules that decrease sterol levels in Niemann-Pick type C mutant cells. PLoS One. 2010;5(9):e12788.

30. Mukherjee S, Maxfield FR. Membrane domains. Annu Rev Cell Dev Biol. 2004;20:839–66.

31. Maxfield FR. Weak bases and ionophores rapidly and reversibly raise the pH of endocytic vesicles in cultured mouse fibroblasts. J Cell Biol. 1982;95(2 Pt 1):676–81.

32. Yamashiro DJ, Fluss SR, Maxfield FR. Acidification of endocytic vesicles by an ATP-dependent proton pump. J Cell Biol. 1983;97(3):929–34.

33. Basu SK, Goldstein JL, Anderson RG, Brown MS. Monensin interrupts the recycling of low density lipoprotein receptors in human fibroblasts. Cell. 1981;24(2):493–502.

34. Lyoumi S, Abitbol M, Rainteau D, Karim Z, Bernex F, Oustric V, et al. Protoporphyrin retention in hepatocytes and Kupffer cells prevents sclerosing cholangitis in erythropoietic protoporphyria mouse model. Gastroenterology. 2011;141(4):1509–19, 19 e1-3.

35. Pipalia NH, Huang A, Ralph H, Rujoi M FR. M. Automated microscopy screening for compounds that partially revert cholesterol accumulation in Niemann-Pick C cells. J Lipid Res 2006;47(2):284–301.

36. Jia Y, Bhuiyan MJ, Jun HJ, Lee JH, Hoang MH, Lee HJ, et al. Ursolic acid is a PPAR-alpha agonist that regulates hepatic lipid metabolism. Bioorg Med Chem Lett. 2011;21(19):5876–80.

37. Peluso MR. Flavonoids attenuate cardiovascular disease, inhibit phosphodiesterase, and modulate lipid homeostasis in adipose tissue and liver. Exp Biol Med (Maywood). 2006;231(8):1287–99.

